# Odor-gated oviposition behavior in an ecological specialist

**DOI:** 10.1101/2022.09.23.509164

**Authors:** Raquel Álvarez-Ocaña, Michael P. Shahandeh, Vijayaditya Ray, Thomas O. Auer, Nicolas Gompel, Richard Benton

## Abstract

Colonization of a novel ecological niche can require, or be driven by, evolution of an animal’s behaviors promoting their reproductive success in the new environment. Little is known about the underlying mechanisms. We have exploited an emerging genetic model for behavioral neuroecology, *Drosophila sechellia* – a close relative of *Drosophila melanogaster* that exhibits extreme specialism for *Morinda citrifolia* noni fruit – to study the evolution and sensory basis of oviposition. *D. sechellia* produces fewer eggs compared to other drosophilids, but lays these almost exclusively on noni substrates, contrasting with avoidance or indifference of noni by generalist species. Visual, textural and social cues do not explain the species-specificity of this preference. By contrast, loss of olfactory input in *D. sechellia*, but not *D. melanogaster*, essentially abolishes egg-laying, suggesting that this sensory modality gates gustatory-driven noni preference. We find the noni bouquet is detected by redundant olfactory pathways. By parsing the fruit’s volatile chemicals and genetic perturbation of individual olfactory pathways in *D. sechellia*, we discover a key role for hexanoic acid and its cognate receptor, the Ionotropic receptor Ir75b, in odor-evoked oviposition. Through receptor exchange in *D. melanogaster*, we provide evidence for a causal contribution of odor-tuning changes in Ir75b to the evolution of oviposition behavior during *D. sechellia*’s host specialization.

## Introduction

Colonization of, and specialization on, a new ecological niche by an animal can provide many benefits, such as access to new resources, protection from biotic and abiotic threats, and avoidance of competition (Sexton et al., 2017). Niche specialization often requires adaptation of multiple behavioral, physiological and morphological traits to survive and reproduce in a new habitat. Divergence of many traits can potentially lead to reproductive isolation and ultimately speciation, making niche specialization a likely driver of biodiversity (Caillaud and Via, 2000; Rundle and Nosil, 2005; Seehausen, 2006). Many striking examples of adaptations to new niches are known, from the rapid evolution of beak morphology of Darwin’s finches as they radiated across the Galápagos archipelago (Grant and Grant, 2005) to visual system loss in Mexican tetra (*Astyanax mexicanus*, blind cave fish) in cave dwellings in the Gulf of Mexico and Rio Grande (Maldonado et al., 2020). While candidate genomic regions and genes have been implicated in some of these adaptations (e.g., (Abzhanov et al., 2004; Lamichhaney et al., 2015)), the restricted genetic tractability of these species – and most other examples in nature – limits our understanding of the mechanistic basis of evolutionary adaptations.

The fly *Drosophila sechellia* provides an exceptional model to investigate the genetic and cellular basis of niche adaptation (Auer et al., 2021; Jones, 2005; Stensmyr, 2009). This species is endemic to the Seychelles archipelago, where it has evolved an extreme specialist lifestyle, feeding and breeding exclusively upon the “noni” fruit of the *Morinda citrifolia* shrub. Adaptation to this niche has occurred in the last few 100,000 years, potentially only since its divergence from a last common ancestor with the cosmopolitan generalist, *Drosophila simulans* (Figure 1A). Importantly, the close phylogenetic proximity of *D. sechellia* to the laboratory model, *Drosophila melanogaster* (Figure 1A), has facilitated the development of genetic tools in this species to explore the mechanistic basis of niche specialization (Auer et al., 2020; Auer et al., 2021; Combs et al., 2018). Previous work has identified *D. sechellia* Odorant receptors (Ors) essential for long-range detection of noni odors, Or22a and Or85c/b (Auer et al., 2020; Dekker et al., 2006; Ibba et al., 2010), and demonstrated a causal relationship between differences in tuning properties of Or22a in *D. melanogaster* and *D. sechellia* and species-specific noni attraction (Auer et al., 2020).

**Figure 1.**
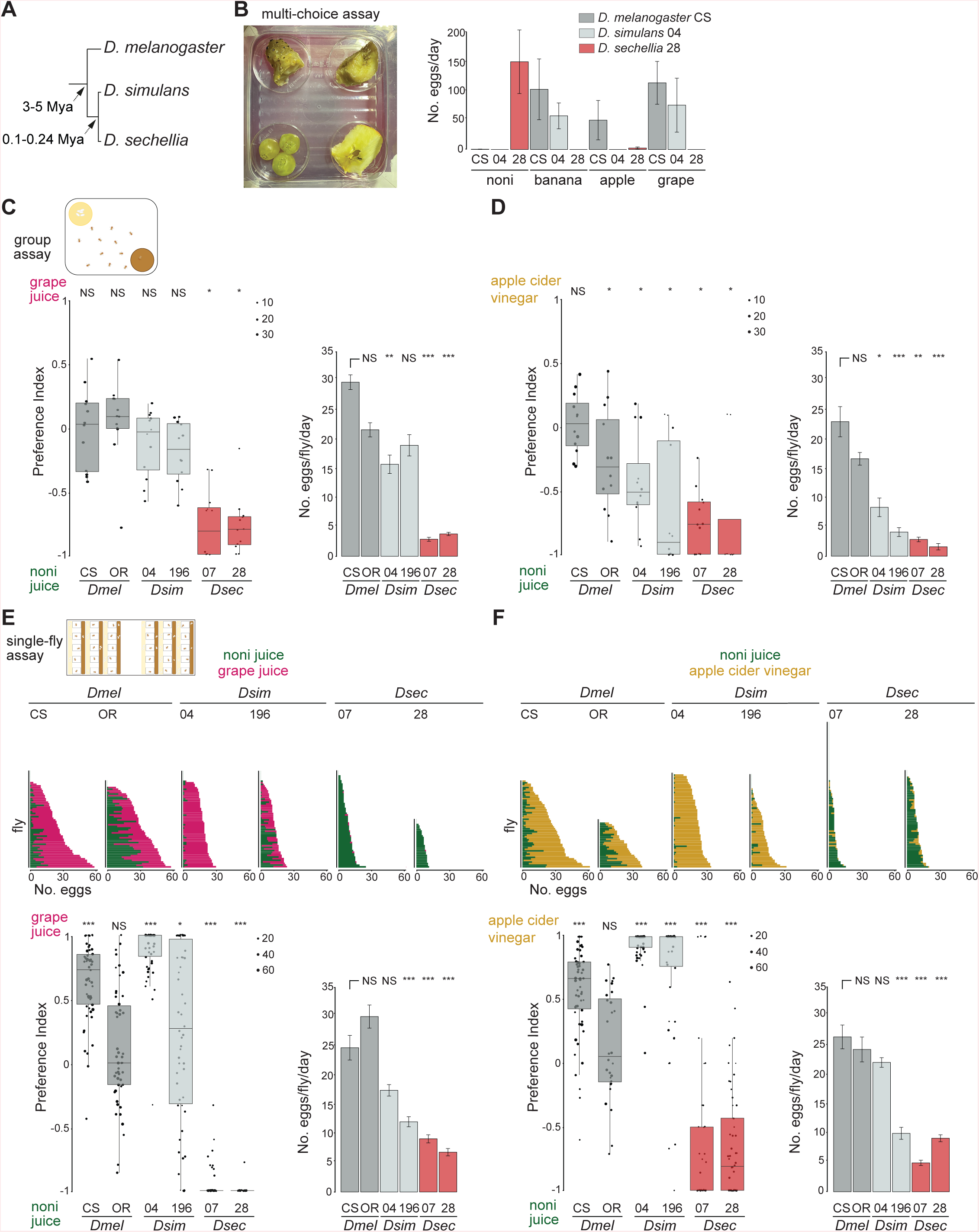
*D. sechellia* displays robust, species-specific preference for oviposition on noni substrates. (A) Phylogeny of the drosophilid species studied in this work. Mya, million years ago. (B) Fruit multiple-choice oviposition preference assay. Left: image of the assay with noni, banana, apple and grape (clockwise from top left) in the arena. Right: quantification of the number of eggs laid per day (n = 3 assays/species, using 50 flies each for a duration of 3 days). Strains used: *D. melanogaster* Canton S (CS), *D. simulans* 14021-0251.004 (04) and *D. sechellia* 14021-0248.28 (28); see Table S1 for details of all strains used in this work. Mean values ± SEM are shown. (C) Group oviposition preference assays for noni juice versus grape juice in 0.67% agarose using two strains each of wild-type *D. melanogaster* (*Dmel*: CS and Oregon R (OR)), *D. simulans* (*Dsim*: 04 and 14021-0251.196 (196)) and *D. sechellia* (*Dsec*: 14021-0248.07 (07) and 28). Left: box plots of oviposition preference index in these assays. In these and all other box plots, the middle line represents the median, and the first and third quartiles correspond to the lower and upper hinges, respectively. Individual data points are overlaid on the box-plots, scaled by the total number of eggs laid in an assay (key at top right of the plot); data beyond the whiskers are considered outliers. For these and other box plots statistical differences from 0 (no preference) are indicated: *** *P* < 0.001; ** *P* < 0.01; * *P* < 0.05; NS (not significant) *P* > 0.05 (Wilcoxon test with Bonferroni correction for multiple comparisons); n = 12 (representing 4 group assays, each scored on 3 successive days with fresh oviposition plates each day). Right: bar plots of egg-laying rate per fly per day in these assays. Mean values ± SEM are shown. Statistically-significant differences from the *D. melanogaster* CS strain are indicated: *** *P* < 0.001; ** *P* < 0.01; * *P* < 0.05; NS *P* > 0.05 (Kruskal-Wallis rank sum test with Nemenyi post-hoc test). (D) Group oviposition preference assays, as in (C), for noni juice versus apple cider vinegar (n = 12, as in (C)). (E) Single-fly oviposition preference assays for noni juice versus grape juice in agarose for the same strains as in (C). Top: total number of eggs laid in each substrate by each female. Bottom left: oviposition preference index. Statistical differences from 0 (no preference) are indicated as in (C). n = 30-60 flies across 1-2 technical replicates. Bottom right: egg-laying rate. Mean values ± SEM are shown; statistical analysis as in (C). (F) Single-fly oviposition preference assays, as in (E), for noni juice versus apple cider vinegar. n = 30-90 flies across 1-3 technical replicates.

Long-range olfactory attraction to noni is only one facet of *D. sechellia*’s phenotypic adaptations in this specialized niche (Auer et al., 2021). Notably, the fruit is highly toxic to other drosophilids (and more divergent insects) – predominantly due to its high levels of octanoic acid – indicating the existence of robust (albeit unclear) resistance mechanisms of *D. sechellia* throughout its life cycle (Legal et al., 1994; Legal et al., 1992; R’Kha et al., 1991). Host fruit toxicity has been suggested to relieve *D. sechellia* from interspecific competition and parasitoidization (Salazar-Jaramillo and Wertheim, 2021), providing a potential explanation for the selective advantage of its stringent niche specialization.

Another unique set of phenotypes of *D. sechellia* relates to the production and deposition of eggs. Compared to its generalist cousins, *D. sechellia* ovaries contain ∼3-fold fewer ovarioles, with a commensurate reduction in egg number (Coyne et al., 1991; Green and Extavour, 2012; R’Kha et al., 1991). The evolutionary advantage (if any) of reduced fecundity is unclear, but may be linked with the larger size of *D. sechellia* eggs (∼50% by volume (Markow et al., 2009)) and the greater tendency of this species to retain fertilized eggs (resulting in facultative ovoviviparity) (Markow et al., 2009). Such observations hint that these traits may be related to more investment of *D. sechellia* in fewer eggs to protect them from the acid-rich noni substrate and/or predators. However, non-adaptive explanations (e.g., pleiotropic effects of mutations in genes underlying other adaptations (Jones, 2004; R’Kha et al., 1997)) cannot be excluded.

Whatever the reason(s) for reduced fecundity of this species, this trait makes the decision of *D. sechellia* females to engage in oviposition particularly important. Previous work has shown that *D. sechellia* depends on the presence of noni for both egg production and laying (Lavista-Llanos et al., 2014; Louis and David, 1986; R’Kha et al., 1991). Furthermore, the most abundant noni chemicals, hexanoic and octanoic acids, can alone induce oviposition (Amlou et al., 1998; Higa and Fuyama, 1993). However, the cognate sensory pathways are unknown, as is the contribution, if any, of other chemosensory or non-chemosensory information to this behavior.

## Results

### Species-specific oviposition preference and rate

To investigate the neurosensory basis of egg-laying behavior in *D. sechellia*, we first compared the specificity of oviposition site selection of *D. sechellia*, *D. simulans* and *D. melanogaster* in a semi-natural, multi-choice assay in which animals were offered slices of different ripe fruits (noni, banana, apple and grape) within an enclosed arena (Figure 1B). While *D. melanogaster* and *D. simulans* flies avoided using noni fruit as an oviposition substrate (preferring two or three of the other fruits instead), *D. sechellia* laid eggs almost exclusively on noni (Figure 1B). To systematically test the species-specificity of oviposition behavior and its sensory basis, we next established two-choice group oviposition assays. In these, egg-laying substrates comprised of commercial juices/vinegar (to ensure consistency of chemical stimulus) mixed in either agarose or Formula 4-24® instant *Drosophila* medium (hereafter, “instant medium”). The latter substrate supported higher egg-laying rate that was important when examining the influence of single odors in subsequent experiments. Two independent strains of each species were tested in all assays to distinguish interspecific from intraspecific differences. *D. melanogaster* and *D. simulans* strains generally exhibited indifference between noni and grape juice substrates, in either agarose or instant medium, with small differences between strains (Figure 1C and Figure S1A). By contrast, *D. sechellia* consistently displayed a strong preference for noni juice-containing substrates (Figure 1C and Figure S1A). Similar, though less marked, differences between species’ preferences were observed in assays offering a choice between noni juice and apple cider vinegar in agarose (Figure 1D), but not in instant medium (Figure S1B).

As social interactions can influence drosophilids’ oviposition preference (Churchill et al., 2021; Elsensohn et al., 2021), we also established a single-fly oviposition assay (see Methods and Figure S2) (Gou et al., 2016). Using the same combinations of stimuli and substrates as in group assays, we observed even more marked species differences in oviposition site preference: *D. sechellia* laid the vast majority of its eggs on noni juice substrates in all assays (Figure 1E-F and Figure S1C-D), while *D. melanogaster* and *D. simulans* exhibited strong preference for either grape juice or apple cider vinegar in agarose, and variable levels of preference for these counter-stimuli in instant medium (with the exception of one strain of *D. simulans*).

Both group and single-fly assays also confirmed the substantially lower fecundity of *D. sechellia* compared to *D. melanogaster* and *D. simulans* (Figure 1C- F and Figure S1). Quantification of eggs laid by individual flies revealed large variation in egg-laying rate for all species, even within a given assay (Figure 1E-F and Figure S1C-D). However, on average, *D. sechellia* consistently laid a low number of eggs (∼5-7 eggs/female/day), while the mean egg-laying frequency for the other species could vary substantially (from ∼10 to ∼30 eggs/female/day), which may be related to the provision of more or less appealing substrates in different assays.

Together, these results highlight the innate, social-context independent, and robust preference for *D. sechellia* for noni substrates, contrasting with the more context-dependent noni indifference or avoidance exhibited by the generalist drosophilids.

### *D. sechellia* exhibits robust probing of the oviposition substrate

In the oviposition experiments with *D. sechellia* on agarose, we observed many small indentations in the substrate surface at the end of the assay (Figure 2A). Such indentations were only occasionally observed on the agarose substrates where *D. melanogaster* or *D. simulans* had laid eggs (data not shown). (The presence of indentations could not be easily assessed in the instant medium substrate due to its more granular texture). Furthermore, indentations were not observed on agarose exposed only to *D. sechellia* males (data not shown), suggesting that they are not the result of non-sexually dimorphic behaviors such as proboscis probing of the substrate.

**Figure 2.**
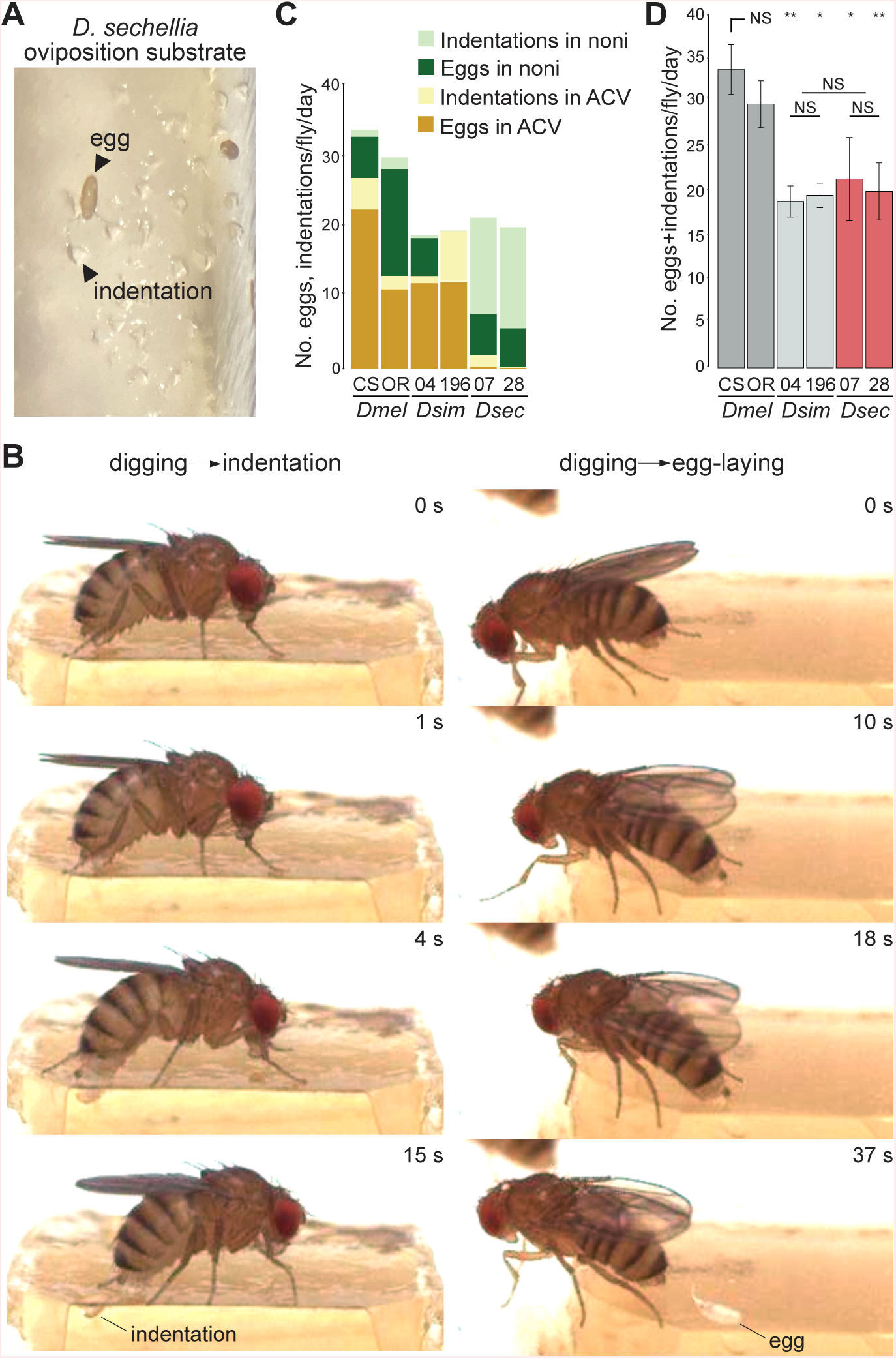
*D. sechellia* makes frequent substrate indentations during oviposition. (A) Photo of the noni juice/agarose substrate at the end of a single-fly oviposition assay with *D. sechellia* illustrating the many indentations in the agarose surface and rare eggs. (B) Still images from high-speed video sequences of *D. sechellia* oviposition behavior illustrating a digging event that does not lead to egg deposition, which results in the formation of a visible indentation on the substrate (left), and digging event that culminates in egg deposition (right). The full videos are provided in Videos S4 and S5. (C) Quantification of the number of eggs and indentations produced by different species and strain on different substrates in 19-28 single-fly two-choice oviposition assays with noni juice and apple cider vinegar (ACV). (D) Rate of summed egg-laying and indentations events observed in the experiments in (C). Mean values ± SEM are shown. Statistically-significant differences from the CS strain are indicated: ** *P* < 0.01; * *P* < 0.05; NS *P* > 0.05 (Kruskal-Wallis rank sum test with Nemenyi post-hoc test).

The size and shape of the indentations led us to hypothesize that they correspond to the substrate marks formed by the ovipositor during “burrowing” – rhythmic digging of the ovipositor into the substrate prior to egg deposition – described in *D. melanogaster* (Cury and Axel, 2021). To test this hypothesis, we used high-speed imaging to visualize *D. sechellia* oviposition behavior at high spatio-temporal resolution (Bracker et al., 2019). Notably, although the total number of egg-laying events captured was low (precluding detailed quantitative analyses), we observed frequent interactions of the ovipositor with the substrate that did not culminate in egg deposition. These interactions ranged from simple substrate touching or scratching by the ovipositor (Video S1-2; see Methods for classification of behaviors) to more involved digging behaviors (Video S3). In two instances, such digging resulted in the formation of a visible indentation (Figure 2B and Video S4). However, post-hoc observation of the substrate revealed several other examples of indentations that were not captured during the recordings, possibly because they were not visible at the camera angle to the substrate. Conversely, we did not observe any other behaviors of the fly that could explain the formation of the indentations. Very similar ovipositor digging events were observed prior to egg laying (Video S5).

These observations support the hypothesis that indentations represent aborted oviposition events in *D. sechellia*. We reasoned that they provide a relevant complementary measure of oviposition behavior to the numbers of eggs laid. We therefore quantified the number of indentations and eggs for all three species in single-fly two-choice assays (Figure 2C). The total number of indentations and eggs were comparable for *D. sechellia* and *D. simulans* strains, and slightly lower than for *D. melanogaster* (Figure 2D). These observations indicate that, despite the much lower egg number than the generalist species, *D. sechellia* still robustly probes the oviposition substrate (see Discussion).

### No evidence for contribution of visual cues to *D. sechellia*’s species-specific oviposition preference

To assess the sensory basis of *D. sechellia*’s strong preference for oviposition on noni substrates, we first tested the contribution of vision as the natural fruits (as well as the artificial substrates) have characteristic colors that might influence decisions on where to lay eggs. In single-fly two-choice assays run in the dark*, D. sechellia* retained very strong, species-specific, preference for laying on noni juice substrates, and no decrease in egg-laying rate was noted (Figure 3A). We extended this analysis to examine whether *D. sechellia* exhibits any unique color preference, reflecting its preference for ripe (dull white/yellow) over unripe (green) fruit. As the noni juice colored the oviposition substrate brown, we tested this possibility using a short-range trap assay (Prieto-Godino et al., 2017), in which identical odor traps – containing noni juice for *D. sechellia* or balsamic vinegar for *D. melanogaster* and *D. simulans* – were enclosed within green or white casings (Figure 3B). Because noni fruit may be found among green foliage, or on the white sandy substrate below *Morinda citrifolia* shrubs (Auer et al., 2021), we reasoned that color contrast may also play an important role in substrate preference, and therefore tested trap preference on a white or green background, as well as in the dark as a control. We observed no preference of any species to enter different colored traps (Figure 3B), suggesting that color is not a critical cue that *D. sechellia* uses to locate host fruit, at least at short-range.

**Figure 3.**
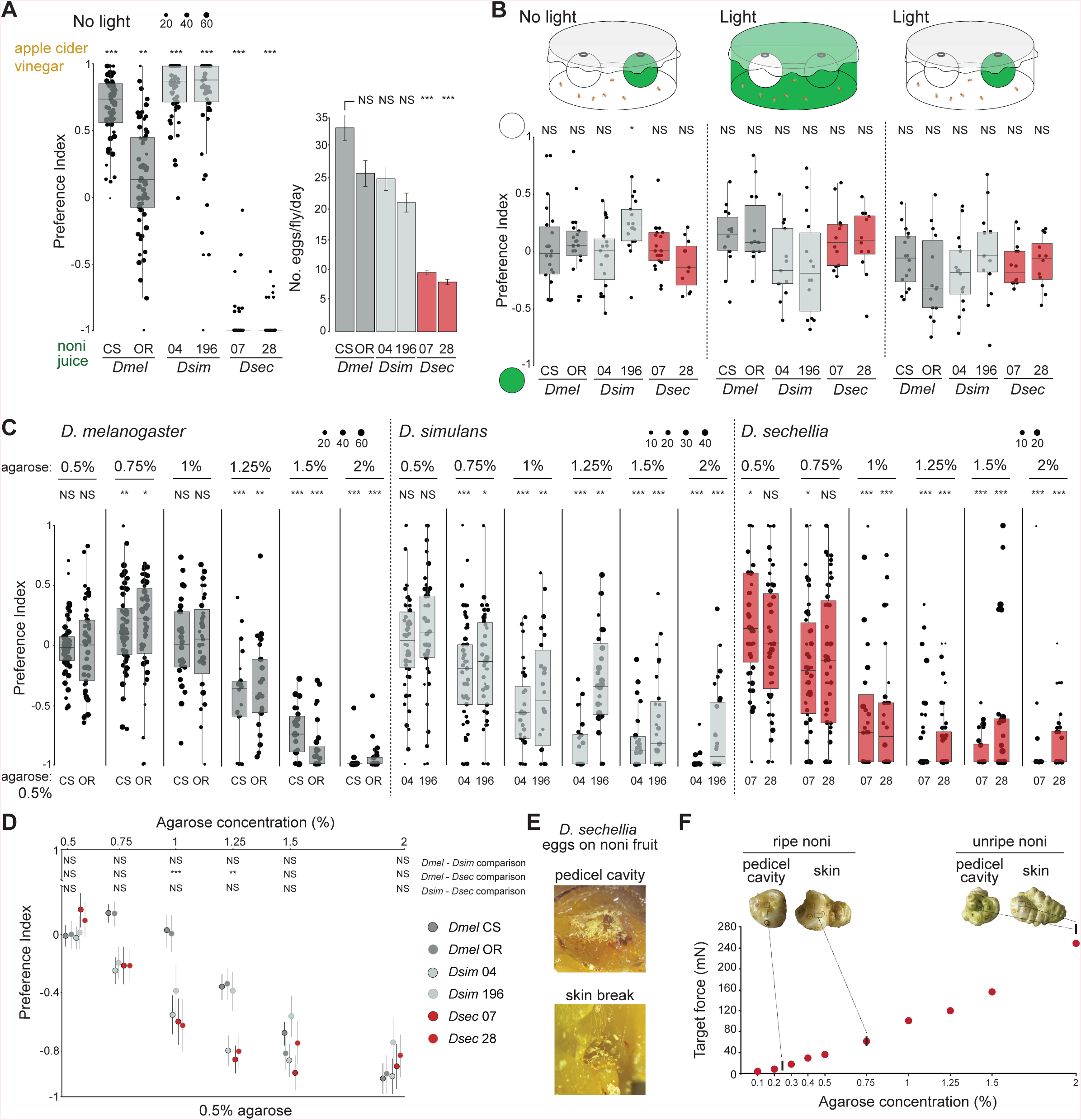
Analysis of visual and textural contributions to *D. sechellia*’s noni preference. (A) Single-fly oviposition preference assays in the dark for noni juice versus apple cider vinegar in agarose (fly strains as in Figure 1C). Left: oviposition preference index. Statistical differences from 0 (no preference) are indicated: *** *P* < 0.001; ** *P* < 0.01; NS *P* > 0.05 (Wilcoxon test with Bonferroni correction for multiple comparisons); n = 60 flies across 2 technical replicates. Right: egg-laying rate in these assays. Mean values ± SEM are shown (Kruskal-Wallis rank sum test with Nemenyi post-hoc test). (B) Group color preference assays in which flies are given a choice to enter two traps containing the same chemical stimulus (balsamic vinegar (*D. melanogaster* and *D. simulans*) or noni juice (*D. sechellia*)) and distinguished only by colored casings with different light and background conditions. Statistical differences from 0 (no preference) are indicated: * *P* < 0.05; NS *P* > 0.05 (Wilcoxon test with Bonferroni correction for multiple comparisons); n = 12-24 assays across at least 2 technical replicates. (C) Single-fly oviposition preference assays testing between the indicate agarose concentrations on the top and 0.5% agarose in the counter-substrate. Both substrates contain apple cider vinegar (*D. melanogaster* and *D. simulans*) or noni juice (*D. sechellia*). Statistical differences from 0 (no preference) are indicated: *** *P* < 0.001; **P < 0.01 * *P* < 0.05; NS *P* > 0.05 (Wilcoxon test with Bonferroni correction for multiple comparisons); n = 30-60 flies across 1-2 technical replicates. (D) Graph recapitulating data from (C). Dots represent the mean values and the bars represent ±SEM. The statistical values represent the most similar strains of the different species: *** *P* < 0.001; ** *P* < 0.01; NS *P* > 0.05 (Kruskal-Wallis rank sum test with Nemenyi post-hoc test). (E) Close-up image of noni fruit illustrating the concentration of *D. sechellia* eggs in the pedicel cavity (where the fruit was attached to the stem) and in the flesh exposed by a skin break. Flies were placed in a group assay oviposition chamber containing whole noni fruits during 72 h. (F) Graph of stiffness of substrates of different agarose concentrations (in noni juice), overlaid with the stiffness ranges of unripe and ripe noni fruits (illustrated in the photos) within the pedicel cavity or on the external skin. Measurements were made using Semmes-Weinstein Monofilaments following the procedure described in (Sanchez-Alcaniz et al., 2017).

### D. sechellia and D. simulans prefer softer substrates compared to D. melanogaster

Substrate hardness is another potential factor influencing oviposition site preference that might have diverged between species. For example, the pest species *D. suzukii* – which oviposits in various ripe, but not rotten, fruits – exhibits stronger preference for stiffer substrates (that presumably resemble more closely ripe fruit) than *D. melanogaster* (Karageorgi et al., 2017). We compared the texture preference profile of *D. sechellia*, *D. simulans* and *D. melanogaster* through single-fly two-choice assays in which both substrates contain the same attractive chemical stimulus (either noni juice or apple cider vinegar) but different stiffness, obtained by pairing a soft agarose substrate (0.5%) with one ranging from 0.5-2% agarose. Although all three species preferred to oviposit on softer agarose, the discrimination threshold was different: *D. melanogaster* only exhibited such a preference when 0.5% was paired with 1.25% (or higher) agarose, while *D. simulans* and *D. sechellia* discriminated a more subtle difference in texture, preferring 0.5% agarose over 0.75% (Figure 3C-D). Textural discrimination ability of *D. sechellia* therefore cannot explain its ecological specialization. However it is consistent with our observations that on native fruits, *D. sechellia* lays its eggs on the softest part of the fruit: the pedicel cavity in intact fruits (or internal flesh in cut/broken fruits) (Figure 3E), which is much softer that the fruit skin, whose stiffness is approximately equivalent to 0.75% agarose (Figure 3F).

### Olfactory pathways required for oviposition and oviposition preference

Having excluded the importance of vision for *D. sechellia*’s egg-laying preference (Figure 3A-B), we reasoned that olfactory cues are likely to be the first sensory signals that *D. sechellia* uses when assessing potential oviposition sites, as these do not require direct contact with the substrate. We first tested near-anosmic double mutant animals for the conserved olfactory co-receptors Orco (required for the function of all Ors) and Ir8a (required for the function of volatile acid-sensing Irs) (Abuin et al., 2011; Auer et al., 2020; Benton et al., 2006; Larsson et al., 2004). Strikingly, in single-fly assays – offering a choice of noni juice and apple cider vinegar in agarose – these flies laid essentially no eggs (Figure 4A-B). Moreover, no indentations were observed on the substrate at the end of the assay (Figure 4B). This lack of oviposition activity is not due to any overt locomotor defects, as these mutant animals display similar levels of activity as wild-type strains (Figure S3). It is also not due to any decrease in egg production, as their ovaries contain a similar number of mature eggs as in wild-type animals (Figure 4C). Importantly, equivalent *D. melanogaster* near-anosmic *Ir8a^1^,Orco^1^* double-mutant animals lay many eggs (Figure S4). These observations provide evidence that olfactory input is critical for oviposition behavior in *D. sechellia*, but not *D. melanogaster*.

**Figure 4.**
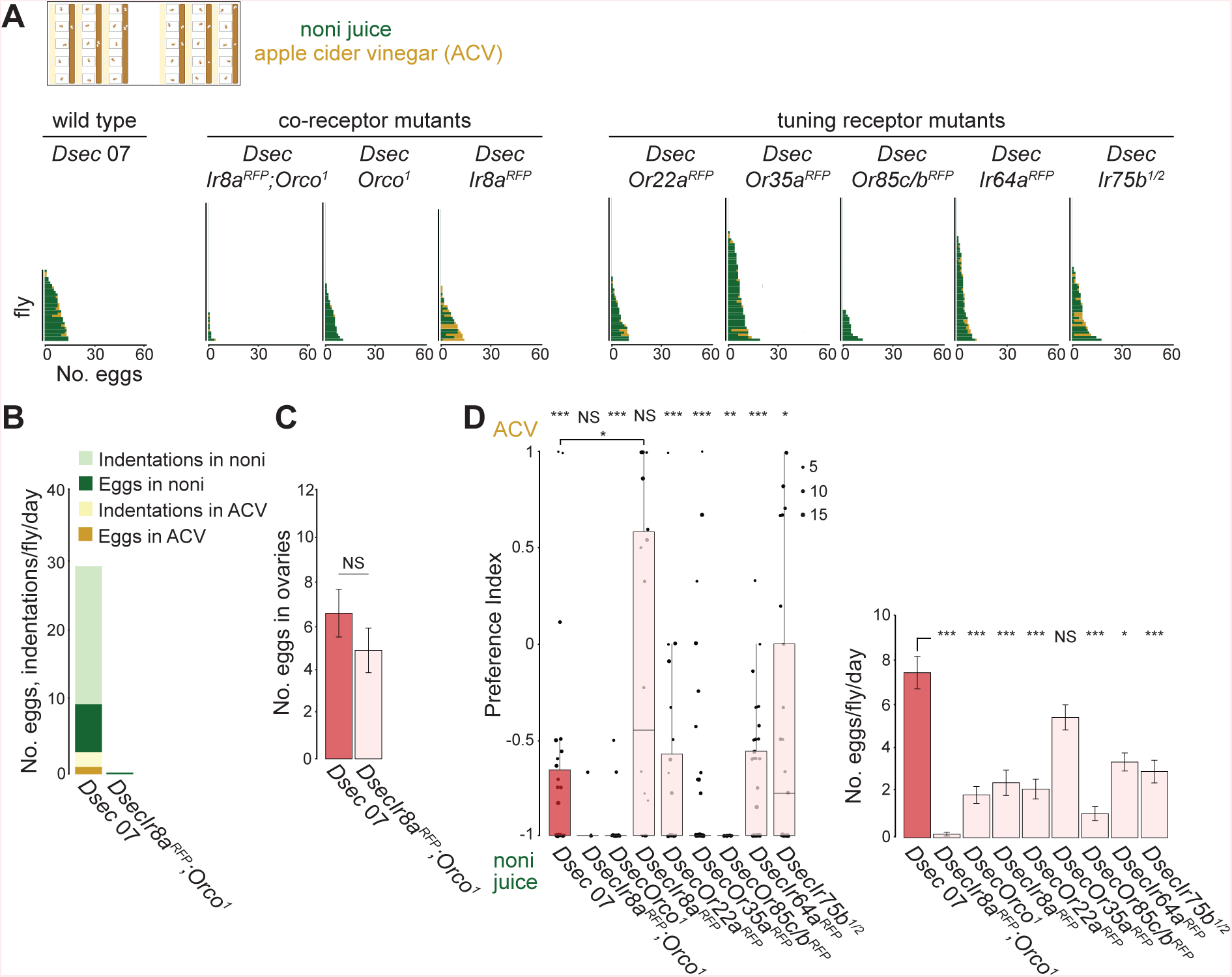
Olfactory pathways required for *D. sechellia* oviposition. (A) Single-fly oviposition preference assays for noni juice versus apple cider vinegar in agarose for the indicated genotypes (Table S1). The plots show the number of eggs laid per fly (n = 30-60 flies across 1-2 technical replicates). *DsecIr75b^1/2^* is a transheterozygous mutant combination. (B) Quantification of the number of eggs and indentations on different substrates of the indicated genotypes (n = 30 (*Dsec* 07) and 58 (*DsecIr8a^RFP^,Orco^1^*) across 1- 2 technical replicates). (C) Mean number of mature eggs per fly (i.e., a pair of ovaries) of the indicated genotypes. Mean values ± SEM are shown. NS *P* > 0.05 (two-sample t-test); n = 9-10 flies. (D) Left: oviposition preference index for the assays shown in (A). Statistical differences from 0 (no preference) are indicated: *** *P* < 0.001; ** *P* < 0.01; * *P* < 0.05; NS *P* > 0.05 (Wilcoxon test with Bonferroni correction for multiple comparisons); n = 30-60 flies across 1-2 technical replicates. *Dsec* 07 and *DsecIr8a^RFP^* show statistical difference *(P =* 0.0328; Wilcoxon test with Bonferroni adjustment). Right: egg-laying rate. Mean values ± SEM are shown. Statistically- significant differences from the *Dsec* 07 strain are indicated: *** *P* < 0.001; * *P* < 0.05; NS *P* > 0.05 (Kruskal-Wallis rank sum test with Nemenyi post-hoc test). The non-significant PI for the *DsecIr8a^RFP^,Orco^1^* double mutant was calculated from the 4/60 animals that laid >2 eggs.

To test whether olfactory cues are sufficient to promote oviposition, we performed a no-choice oviposition assay in which flies were provided with an agarose/sucrose substrate with a non-accessible source of noni juice or, as control, water (Figure S5). No differences were observed in egg-laying rate of *D. sechellia* (or *D. melanogaster*) strains between these substrates (Figure S5). These results argue that noni odors alone are insufficient to promote egg-laying behavior, which presumably relies also upon gustatory input through multiple contact chemosensory organs, as is the case in *D. melanogaster* (Chen et al., 2022; Chen and Amrein, 2017; Chen and Dahanukar, 2020).

To further understand the contribution of olfaction to *D. sechellia*’s oviposition behavior, we next tested the *Orco* and *Ir8a* co-receptor mutants singly: both showed a decreased egg-laying rate compared with wild-type controls, but only *Ir8a* mutants displayed reduced preference for noni juice (Figure 4A,4D). Similar phenotypes for these mutants were observed in two-choice assays with grape juice as a counter-stimulus (Figure S6). We went on to screen the phenotypes of mutants lacking genes encoding individual odor-specific “tuning” Ors and Irs (Auer et al., 2020), including Or22a and Or85c/b, which are required for long-range noni attraction (Auer et al., 2020). While these lines displayed variable reductions in egg-laying rate, none of them displayed significantly diminished oviposition preference for noni substrates (Figure 4A,4D and Figure S6). The maintenance of robust oviposition preference towards noni in most of these assays suggested that multiple, partially redundant olfactory signals contribute to oviposition behavior in *D. sechellia*.

### Analysis of the effect of individual noni chemicals on oviposition

To characterize the noni chemicals promoting *D. sechellia*-specific oviposition, we tested several candidates in single-fly assays using the different species and three odor concentrations (0.05%, 0.1%, 0.5%) (Figure 5 and Figure S7). Confirming and extending previous group assays (Amlou et al., 1998; Higa and Fuyama, 1993; Matsuo et al., 2007), we found that hexanoic acid promoted very strong preference in *D. sechellia* at all concentrations tested and a higher egg-laying rate (compared to water-only control substrates) at least at intermediate concentrations (Figure 5A- B). *D. melanogaster* and *D. simulans* both display slight preference or indifference at lower concentrations of hexanoic acid and strong aversion at the highest concentration (Figure 5A-B). Octanoic acid has also been described to be an oviposition stimulant/attractant for *D. sechellia* in some (Amlou et al., 1998; Legal et al., 1999; Matsuo et al., 2007), though not all (Markow et al., 2009) reports. In our assays, this acid did not evoke strong oviposition preference of *D. sechellia* at lower concentrations; moreover, egg-laying was largely suppressed at the highest concentration, although the very few eggs laid were found on the octanoic acid substrate (Figure 5A-B). *D. melanogaster* and *D. simulans* found this odor generally aversive (Figure 5A).

**Figure 5.**
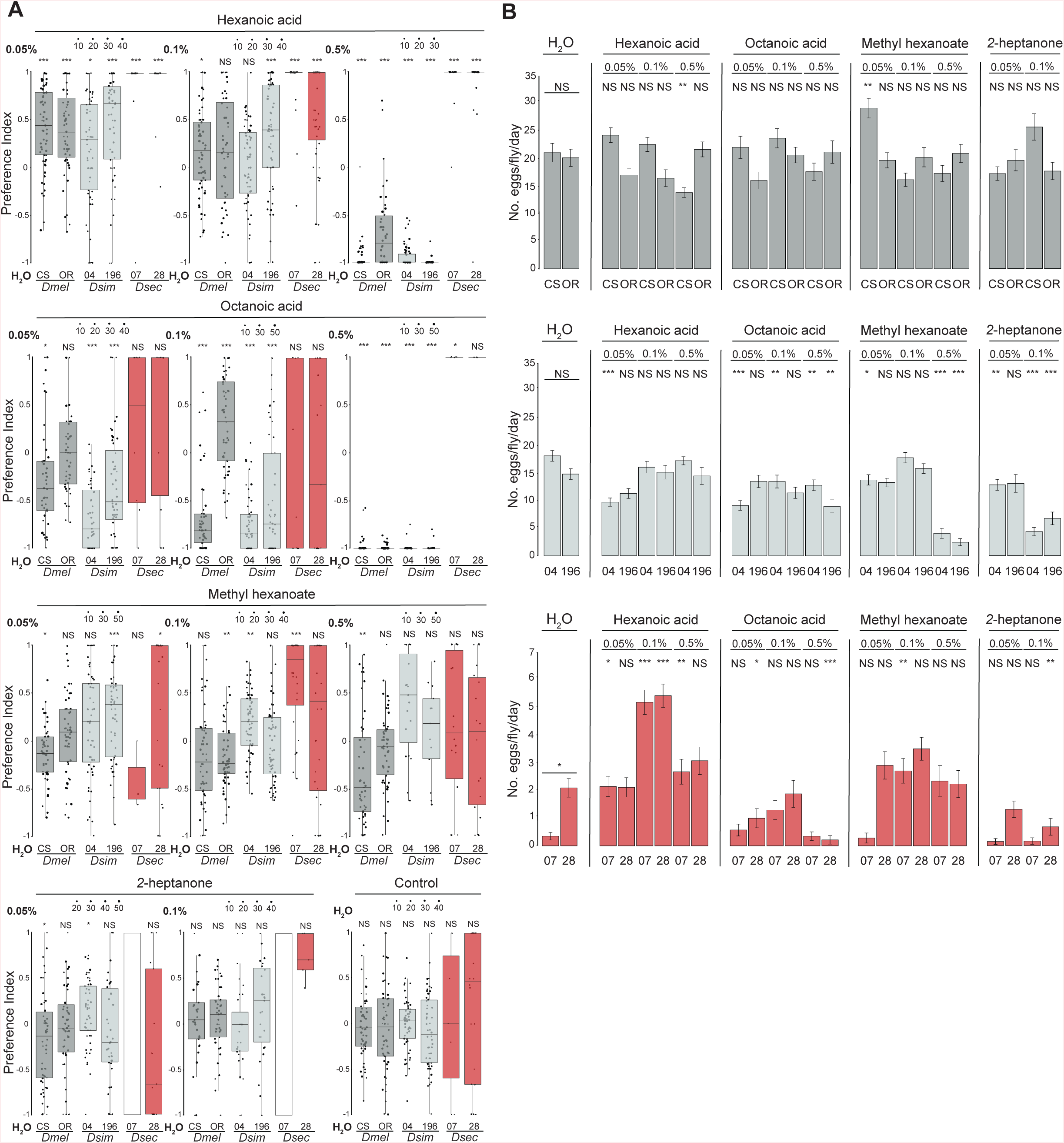
Analysis of the effect of individual noni chemicals on oviposition. (A) Single-fly oviposition assays of the indicated strains testing different odors and concentrations in an instant medium substrate. Oviposition preference index. Statistical differences from 0 (no preference) are indicated: *** *P* < 0.001; ** *P* < 0.01; * *P* < 0.05; NS *P* > 0.05 (Wilcoxon test with Bonferroni correction for multiple comparisons); n = 30-60 flies across 1-2 technical replicates. For *Dsec* 07 assays with *2-*heptanone (indicated with a white rectangle), the low number of flies laying eggs prevented calculation of a preference index. (B) Egg-laying rate of the assays in (A). Mean values ± SEM are shown. Statistical comparisons of the effect of odors on egg-laying rate were performed across strains: *** *P* < 0.001; ** *P* < 0.01; * *P* < 0.05; NS *P* > 0.05 (Kruskal-Wallis rank sum test with Nemenyi post-hoc test).

We tested two other noni chemicals that are behaviorally-important for long-range noni location: methyl hexanoate (detected by Or22a) and *2*-heptanone (detected by Or85c/b) (Auer et al., 2020; Dekker et al., 2006; Ibba et al., 2010). Methyl hexanoate stimulated a slight enhancement of egg-laying at intermediate concentrations, but flies did not display a strong oviposition site preference for substrates containing this chemical (Figure 5A-B). Similarly, neither *D. melanogaster* nor *D. simulans* exhibited strong preference or aversion to methyl hexanoate-containing substrates. *2*-heptanone had little influence on oviposition-site selection of any species, and was highly toxic for flies at the highest (0.5%) concentration (Figure 5A and data not shown).

Lastly, we tested oviposition stimulants described in *D. melanogaster*, valencene and limonene, which are detected by Or19a neurons (Dweck et al., 2013). *D. sechellia* flies were indifferent to or avoided oviposition on substrates containing either of these chemicals, and egg-laying was suppressed at high stimulus concentrations (Figure S7). Interpretation of these results must be tempered, however, with our inability to consistently reproduce behavioral effects of these compounds on oviposition reported in *D. melanogaster* (Figure S7) (Dweck et al., 2013), potentially reflecting observations that the behavioral function of this olfactory pathway is context dependent (Chin et al., 2018).

Together these experiments reveal the complex, concentration-dependent influence of different individual chemicals on drosophilid oviposition behavior, which might be due to their detection via both olfactory and gustatory systems. Nevertheless, the robust and *D. sechellia*-specific effect of hexanoic acid on oviposition led us to focus on determining the sensory mechanism by which this noni chemical is detected.

### Olfactory detection of hexanoic acid by Ir75b promotes oviposition in *D. sechellia*

To define the sensory mechanisms of volatile hexanoic acid-mediated control of oviposition behavior, we tested our panel of *D. sechellia* olfactory receptor mutants. Loss of either Ir8a or Orco alone led to abolished or greatly diminished egg-laying on hexanoic acid substrates (Figure 6A), suggesting that both Ir and Or pathways contribute. Within the Ir repertoire, Ir75b was an excellent candidate as this receptor has evolved novel sensitivity to hexanoic acid in *D. sechellia* from the ancestral butyric acid sensitivity of the *D. melanogaster* and *D. simulans* orthologs (Prieto-Godino et al., 2017). Indeed, mutation of *Ir75b* in *D. sechellia* led to complete loss of egg-laying on hexanoic acid substrates, a phenotype confirmed in two independent alleles, and a transheterozygous *Ir75b* mutant combination (Figure 6A). Dissection of these flies’ ovaries revealed a similar number of eggs as in controls (Figure 6B), suggesting the defect was in egg-laying not production. Consistent with this hypothesis, *DsecIr75b* mutant flies also produced no indentations in these assays (Figure 6C). By contrast, *D. sechellia* lacking the broadly-tuned acid-sensor, Ir64a (Ai et al., 2010), still oviposited, laying almost all eggs on hexanoic acid substrates (Figure 6A).

**Figure 6.**
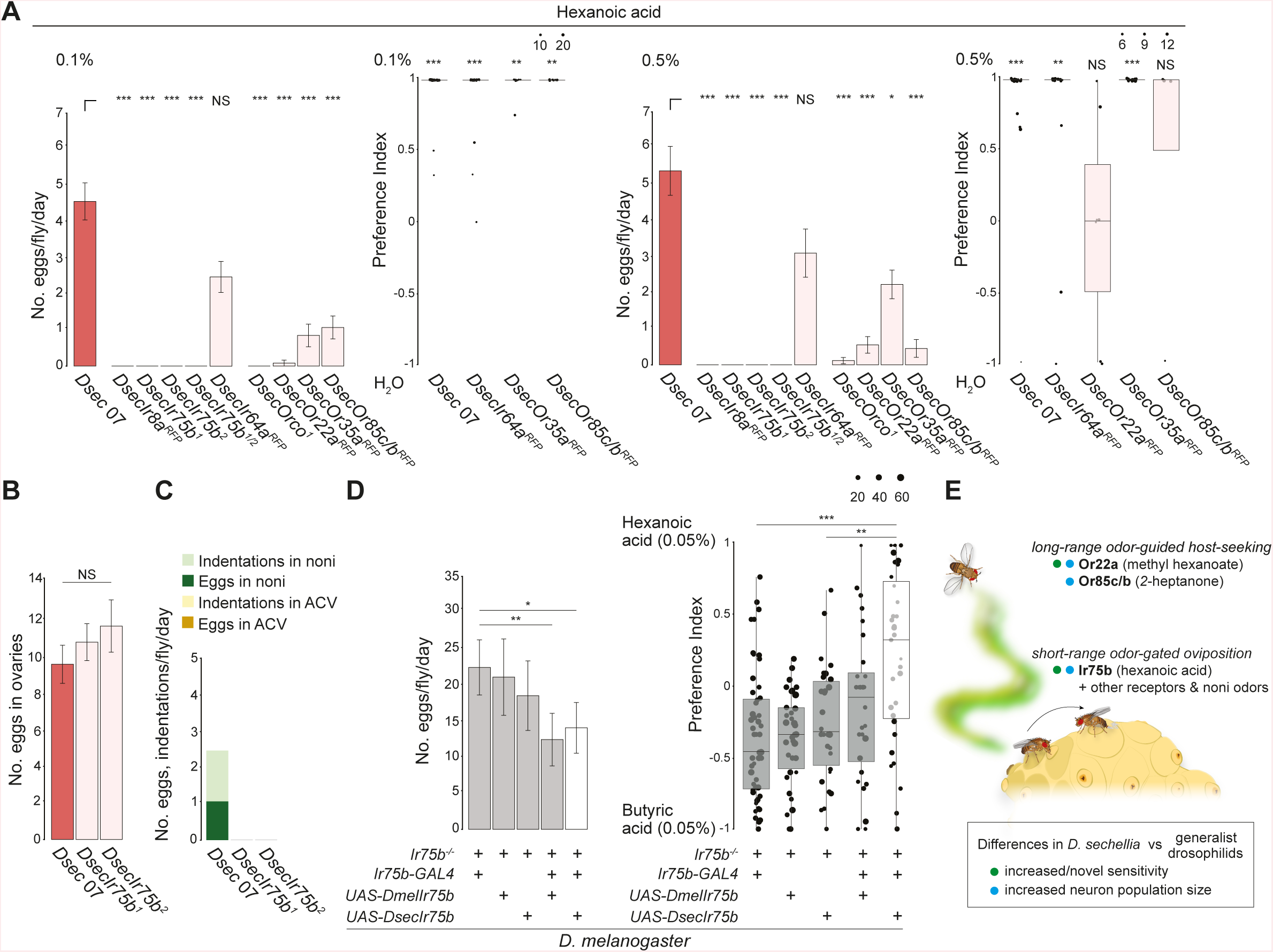
*D. sechellia* Ir75b is required for hexanoic acid responses and sufficient to shift oviposition preference in *D. melanogaster*. (A) Single-fly oviposition assays testing H_2_O versus 0.1% hexanoic acid and H_2_O versus 0.5% hexanoic acid in instant medium. Left: oviposition preference indices are only shown for genotypes that laid 2 or more eggs in these assays. Statistical differences from 0 (no preference) are indicated: *** *P* < 0.001; ** *P* < 0.01; NS *P* > 0.05 (Wilcoxon test with Bonferroni correction for multiple comparisons); n = 30- 83 flies, 1-3 assays. Right: egg-laying rate. Mean values ± SEM are shown. *** *P* < 0.001; * *P* < 0.05; NS *P* > 0.05 (Kruskal-Wallis rank sum test with Nemenyi post- hoc test). (B) Average number of mature eggs per pair of ovaries per fly. Mean values ± SEM are shown. NS *P* > 0.05 (two-sample t-test); n = 9 flies. (C) Quantification of the number of eggs and indentations on different substrates of the indicated genotypes (n = 30 flies in one technical replicate). (D) Single-fly oviposition assays testing 0.05% hexanoic acid versus 0.05% butyric acid of *D. melanogaster Ir75b* mutant and rescue genotypes. Left: egg-laying rate for *Ir75b-Gal4* control (*w;Ir75b-Gal4/+;Ir75b^DsRed^*), *UAS-DmelIr75b* control (*w;UAS- DmelIr75b/+;Ir75b^DsRed^*), *UAS-DsecIr75b* control (*w;UAS-DsecIr75b/+;Ir75b^DsRed^*), *DmelIr75b* rescue (*w;Ir75b-Gal4/UAS-DmelIr75b;Ir75b^DsRed^*), and *DsecIr75b* rescue (*w;Ir75b-Gal4/UAS-DsecIr75b;Ir75b^DsRed^*). *DmelIr75b* and *DsecIr75b* rescue strains showed a significant reduction in the number of eggs compared to *DmelIr75b-Gal4* control. Right: oviposition preference indices for these genotypes. No significant differences were detected between *Ir75b-Gal4* control, *UAS- DmelIr75b* control and *DmelIr75b* rescue strains. The *DsecIr75b* rescue strain showed a significant shift in preference toward 0.05% hexanoic acid compared to both *DmelIr75b-Gal4* and *UAS-DsecIr75b* controls. *** *P* < 0.001; ** *P* < 0.01; *P* < 0.05 (Wilcoxon tests with Bonferroni correction for multiple comparisons). N = 28- 55 flies per genotype measured across at least 2 technical replicates. (E) Schematic summarizing the contributions of different olfactory pathways to niche specialization in *D. sechellia*.

Amongst the Ors, all three mutants (*DsecOr22a*, *DsecOr35a* and *DsecOr85c/b*) displayed reduced egg-laying rate (Figure 6A). Of the subset of flies that did lay eggs, the *DsecOr35a* and *DsecOr85c/b* mutants maintained strong preference for oviposition on hexanoic acid substrates while *DsecOr22a* mutants no longer discriminated this substrate from the control medium (Figure 6A). Or22a neurons are generally considered to be ester sensors in drosophilids (de Bruyne et al., 2010), but weak Or22a-dependent hexanoic responses have been described in *D. sechellia* (Auer et al., 2020) as well as in *D. melanogaster* (Hallem and Carlson, 2006) (where it is the most sensitive hexanoic acid sensor of this species (Munch and Galizia, 2016)), suggesting that it might be a second olfactory pathway for this oviposition stimulant (see Discussion).

### Evolution of Ir75b tuning can explain species-specific behavioral responses

Given the important role of Ir75b for hexanoic acid-stimulated oviposition of *D. sechellia*, we asked if the evolution of the tuning of this receptor might explain species-specific oviposition behavior. In an *Ir75b* mutant of *D. melanogaster* (Mika et al., 2021), we rescued Ir75b function through transgenic expression of either *D. sechellia* Ir75b or, as a control, *D. melanogaster* Ir75b. These flies were offered a choice to lay eggs on substrates containing hexanoic acid or butyric acid, the preferential ligand of the receptors from *D. sechellia* and *D. melanogaster*, respectively (Prieto-Godino et al., 2017). Egg-laying rate was broadly comparable between all control mutant and both rescue lines (Figure 6D). This result indicates that a functional Ir75b pathway is not important for egg-laying in *D. melanogaster* – consistent with our observation that near-anosmic *D. melanogaster* lay many eggs – thereby permitting assessment of the contribution of the Ir75b pathway to oviposition preference. Rescue flies expressing *D. melanogaster* Ir75b displayed a substrate preference that was not significantly different from parental genotypes (Figure 6D). By contrast, expression of *D. sechellia* Ir75b was sufficient to shift oviposition preference from butyric acid to hexanoic acid substrates compared to controls (Figure 6D). These results are consistent with a causal contribution of *Ir75b* to the evolution of oviposition site preference during *D. sechellia*’s host specialization.

## Discussion

Decisions on when and where to lay an egg are critical for all oviparous animals to maximize the chance of survival of their offspring, in particular those lacking parental care (Cury et al., 2019; Rudolf and Rodel, 2005). As such, these decisions are influenced by multiple biotic and abiotic factors in the environment. When species establish themselves within a new ecological niche, changes in these factors can exert selective pressures for novel or modified behavioral responses to sensory cues. It is also possible that chance evolution of traits can permit exploitation of a new niche. Either way, studying such differences between species can provide insight into the relative importance of the plethora of environmental signals, as well as the mechanisms by which nervous systems evolve, changing the relationship between these signals and behavioral outputs. *D. sechellia* offers an excellent opportunity to study oviposition behavioral adaptations, both because its specialist lifestyle likely constrains the set of pertinent sensory cues and because its low fecundity presumably renders the decision to lay an individual egg more important than for highly fertile species. Moreover, the phylogenetic proximity of *D. sechellia* to the generalists *D. melanogaster* and *D. simulans* facilitates comparative behavioral and genetic analyses that might enable reconstruction of the (still-unknown) evolutionary history of this species (Auer et al., 2021; Matsuo, 2008).

Studies in *D. melanogaster* have revealed that oviposition decisions are complex, multisensory-guided behaviors (Cury et al., 2019), and the ultimate choice of egg-laying site is often assay-dependent (e.g., (Schwartz et al., 2012; Yang et al., 2008)). Using several types of behavioral assays, we have confirmed the importance of noni for *D. sechellia* for egg-laying rate and site selection. The latter trait contrasts with the variable preferences of *D. melanogaster* and *D. simulans*. We also discovered an unappreciated feature of oviposition behavior of *D. sechellia*: extensive probing of the substrate surface, resulting in the formation of numerous indentations. These indentations are most likely equivalent to the “burrows” resulting from aborted oviposition events of *D. melanogaster* (Cury and Axel, 2021). One explanation for the high rate of indentations in *D. sechellia* is that females engage in the initiation of the oviposition routine unaware of the low number of eggs they carry. This seems unlikely, however, as it would represent a futile energetic investment for these animals, and *D. melanogaster* mutants that lack eggs do not make indentations (R.A.-O. and R.B., unpublished). Moreover, high-resolution behavioral observations suggest that the presence of the egg in the ovipositor is integral to penetration of the substrate in *D. melanogaster* (Cury and Axel, 2021) and *D. sechellia*. We favor a hypothesis that extensive indentation formation by *D. sechellia* reflects greater choosiness of this species to deposit eggs only after the female has ascertained to have found the optimal substrate available.

To account for the species-specificity of *D. sechellia* substrate selection, we have been able to exclude several sources of sensory information. Visual input is unimportant (at least at short-range), and *D. sechellia* does not exhibit obvious changes in preference for colors that mimic the choice this species makes in nature. While *D. sechellia* prefers to lay eggs within the softest part of the fruit, and within softer agar, there is no difference in texture preference compared to *D. simulans*, suggesting that this trait is not a key facet of host adaptation, contrasting with the fresh-fruit feeder *D. suzukii* (Karageorgi et al., 2017). Finally, although communal egg-laying is widespread in many invertebrates and vertebrates (Doody et al., 2009), we do not find evidence that this phenomenon contributes to noni preference; if anything, isolated flies lay more eggs with stricter noni preference than those in groups.

Our genetic analysis indicates that olfactory input is essential for egg-laying in *D. sechellia*, as near-anosmic flies fail to lay eggs even in the presence of noni despite normal egg production. Conversely, exposure of flies to noni odors alone, without allowing them to have gustatory sensation of noni juice, does not enhance oviposition rate. Together, these observations suggest that both olfactory and gustatory inputs are important: without olfaction, gustatory signals are insufficient for promoting oviposition, but olfactory signals without gustatory inputs are similarly ineffective. A future priority is to determine how *D. sechellia* detects noni via gustation and if, as in the olfactory system, any gustatory pathways differ between drosophilids.

While loss of the vast majority of olfactory input prevents egg-laying on noni in *D. sechellia*, we found substantial redundancy, as loss of any single tuning Or or Ir (or even Orco) did not strongly diminish noni preference. This observation indicates that multiple distinct odors, acting via several different olfactory receptors, must contribute to short-range behavioral decisions. By simplifying the noni odorscape in our oviposition assays we demonstrate the unique oviposition- promoting role of hexanoic acid and, importantly, define Ir75b and its obligate co-receptor Ir8a, as the cognate sensory receptor. Although hexanoic acid might also be detected by gustatory neurons (based upon studies in *D. melanogaster* (Ahn et al., 2017; Sanchez-Alcaniz et al., 2018; Tauber et al., 2017)), the selective expression of Ir75b and Ir8a in the antenna argues that this is an odor-guided behavior. Moreover, the demonstration that Ir75b is required for this behavior provides an explanation for the evolutionary changes described in this sensory pathway: while *D. melanogaster* (and *D. simulans*) Ir75b are tuned primarily to butyric acid, the *D. sechellia* receptor has evolved novel sensitivity to hexanoic acid, through amino acid substitutions within the ligand-binding domain (Prieto-Godino et al., 2017; Prieto-Godino et al., 2021). In addition, *D. sechellia* exhibits a 2-3-fold increase in number of sensory neurons expressing Ir75b, resulting in increased sensory pooling onto partner interneurons in the brain (Prieto-Godino et al., 2017). Importantly, replacement of *D. melanogaster* Ir75b with the *D. sechellia* receptor induces a small but significant shift in oviposition site preference, indicating that receptor tuning changes are sufficient alone to confer more *D. sechellia*-like behavior on *D. melanogaster*.

Together with previous work (Auer et al., 2020), our studies of noni- dependent odor-guided behaviors in *D. sechellia* reveal similarities and differences in the coding and evolution of olfactory pathways mediating long-range and short-range detection (Figure 6E). The high redundancy in short-range olfactory signals contrasts markedly with olfactory contributions to long-range noni host-seeking, where loss of single tuning receptors essentially abolished the ability of flies to locate the odor source (Auer et al., 2020). This difference might reflect the complexity of the noni odor blend at different spatial scales: there are likely to be fewer, highly-volatile, compounds reaching behaviorally-relevant concentrations at a distance compared to odors present at short-range (Auer et al., 2020). Concordantly, the behaviorally most-important receptors for long-range (Or22a and Or85c/b) and short-range (Ir75b) noni detection, display differences in sensitivity: Ir75b neurons require several orders of magnitude higher odor stimulus concentration to evoke the same level of neuronal firing as Or22a or Or85c/b neurons (Auer et al., 2020). The segregation of behavioral function of the pathways is, however, not absolute: the long-range olfactory detectors, notably Or22a, also appear to contribute to oviposition behaviors on hexanoic acid substrates, though further genetic analysis will be necessary in future studies.

One striking commonality of all three of these OSN populations is their expansion in *D. sechellia*, although the functional significance of this phenotype is unknown. By contrast, the nature of odor specificity evolution of these pathways is different: Or85c/b neuron sensitivity to *2*-heptanone is unchanged across the drosophilid species (Auer et al., 2020), *D. sechellia* Or22a has enhanced sensitivity to methyl hexanoate compared to the orthologous receptor in *D. melanogaster* (but not in *D. simulans*) (Auer et al., 2020; Dekker et al., 2006), while Ir75b has acquired new sensitivity to hexanoic acid specifically in *D. sechellia* (Prieto-Godino et al., 2017; Prieto-Godino et al., 2021). In addition, methyl hexanoate and *2*-heptanone, but not hexanoic acid, are emitted by a wide range of fruits (Dweck et al., 2018). A model to unify these observations is that the methyl hexanoate and *2*-heptanone act as “habitat odor cues” (Webster and Carde, 2017), attracting *D. sechellia* (but also other species) in the vicinity of noni, while hexanoic acid is a specific “host odor cue” (Webster and Carde, 2017) that, through Ir75b, evokes short-range behaviors only in *D. sechellia*.

In this context, the ecological role of the Ir75b sensory pathway in *D. melanogaster* is unclear, although optogenetic activation experiments have provided evidence for a role in positional attraction and oviposition preference (Prieto-Godino et al., 2017; Wu et al., 2022). Several other olfactory pathways have been implicated in oviposition promotion in *D. melanogaster* (Cury et al., 2019; Dweck et al., 2015; Lin et al., 2015), including Or19a, which detects the citrus odors valencene and limonene (Dweck et al., 2013). Interestingly, *D. sechellia* Or19a neurons appear to have lost sensitivity to these odors (Dweck et al., 2013), which are not reliably detected in noni fruit (Auer et al., 2020). Adaptation of this species might therefore have involved sensory gain or loss in several olfactory pathways to match the pertinent chemical signals in its niche.

Finally, beyond the issues mentioned above, a key future question – in any species – is how olfactory input controls oviposition behavior. Recent studies in *D. melanogaster* have defined circuitry linking mating and egg-laying (Wang et al., 2020); notably, the activity of some of the component neuron populations (i.e., ovoENs and ovoINs in the central brain) are activated or inhibited by gustatory and mechanosensory input (Wang et al., 2020). It is possible that olfactory sensory pathways (e.g., downstream of the Ir75b sensory input in *D. sechellia*) impinge on this circuitry (Nojima et al., 2021). Alternatively, olfactory signals might have only an indirect influence, for example, by modulating gustatory inputs to this egg-laying circuitry. Further exploration of the neural basis of oviposition in *D. sechellia* should yield insights into the mechanistic basis of the adaptations of this critical behavior to this species’ unique lifestyle.

## Methods

### Drosophila strains

*Drosophila* stocks were cultured in a 25°C incubator under a 12 h light:12 h dark cycle on a standard wheat flour–yeast–fruit juice food. Unless noted otherwise, *D. sechellia* culture vials were supplemented with noni paste, consisting of a few grams of Formula 4-24® instant *Drosophila* medium, blue (Carolina Biological Supply) and noni juice (Raab Vitalfood Bio). All strains used in this study are listed in Table S1 and sources of chemicals are listed in Table S2.

### Oviposition assays

We maximized flies’ egg-laying capacity by following the protocol of (Gou et al., 2016): prior to the experiments, ∼50 1-2 day-old females and males were collected and placed in new fly food tubes enriched with dry yeast (*D. melanogaster* and *D. simulans*) or with dry yeast and noni paste (*D. sechellia*) for 5 days. At this point the food was typically full of crawling larvae, inducing females to retain eggs until transferred to the assay chamber. Unless otherwise stated, oviposition assays were performed at 25°C, 60% relative humidity and a 12 h light:12 h dark cycle (starting assays in the early afternoon), in either an incubator or behavior room for 22-72 h (depending upon the assay, see below).

Previous work suggested that the low egg-laying rate of *D. sechellia* is due to alterations in dopamine metabolism – which contributes, at least indirectly, to fertility in *D. melanogaster* (Gruntenko and Rauschenbach, 2008; Neckameyer, 1996) – and could be partially compensated by supplementation of food with the dopamine precursor 3,4-dihydroxyphenylalanine (L-DOPA), which is found in noni fruit (Lavista-Llanos et al., 2014). To increase *D. sechellia*’s oviposition rate, we cultivated flies’ for five days on noni food supplemented with L-DOPA (1 mg/ml), but did not observe increased egg-laying in either group or single-fly assays compared to control flies given noni paste (Figure S8). Treatment with α-methyl- DOPA (0.4 mM), a non-hydrolysable L-DOPA analog that acts as a competitive inhibitor of DOPA decarboxylase (which converts L-DOPA to dopamine) reduced egg-laying in the single-fly assay but not the group assay (Figure S8). Our inability to fully reproduce the reported effects on oviposition (Lavista-Llanos et al., 2014) – we did not examine other traits investigated in that study, such as egg size and germline cyst apoptosis – might be due to experimental differences in our assays (e.g., use of noni fruit in (Lavista-Llanos et al., 2014)) or the use of more fertile *D. sechellia* strains.

*Fruit multiple-choice assay*: ripe fruits (apple, banana, grape (all from Migros); noni from *M. citrifolia* plants (University of Zurich Botanical Gardens and Canarius) grown in a greenhouse) were cut into thick (1-2 cm) slices. Fruit pieces were placed in a 5 cm Petri dish (Falcon) inside a plastic chamber (15 cm length × 14 cm width × 5 cm height; Migros). For all species, 50 females and 20 males were anesthetized on ice and introduced into the chamber, which was covered with a fabric gauze. The chambers were placed in a behavioral room in constant darkness for 72 h, after which the number of eggs on each fruit piece was quantified.

*Group two-choice assay*: agarose (Promega) substrates were prepared as follows: a 1% agarose solution was prepared and let to cool down until it was possible to hold the glass Erlenmeyer flask with bare hands. The 1% agarose preparation was added to juice/odor solution in a 2:1 ratio, resulting in a final concentration of 0.67% agarose. The final mixture was poured up to a 0.5 cm depth into 3 cm Petri dishes (Falcon). Agarose plates were conserved at 4°C for a maximum of 3 days. Instant medium substrates were prepared by diluting 12 g of instant medium in 100 ml of noni juice (or apple cider vinegar) creating a semi-solid consistency. The instant medium was added into a 3 cm Petri dish until fully covering the bottom of the plate. Instant medium mixes were conserved at 4°C and used within two weeks. Two Petri dishes containing the desired combination of substrates were placed into the same chamber used for the fruit multiple-choice assay. 10 females and 8-10 males (*D. melanogaster* and *D. simulans*) or 20 females and 15-20 males (*D. sechellia*) were anesthetized with CO_2_ and introduced into the chamber for three consecutive days. Due to *D. sechellia*’s low fecundity, through preliminary experiments we considered that the number of eggs laid by 20 *D. sechellia* female flies was sufficient to observe a clear behavioral preference between two conditions. Petri dishes were exchanged with fresh ones every 24 h by quickly lifting the mesh cover. The number of eggs laid per substrate was counted independently on each plate.

*Group no-choice odor cue assay*: as for the “*Group two-choice assay*” except using a single 3 cm diameter Petri dish containing 0.67% agarose and 150 mM sucrose (Sigma) onto which a non-accessible container covered with a fabric gauze and a perforated cap of a 15 ml Falcon tube (diameter 1 cm, height 2 cm; Techno Plastic Products AG) into which 300 μl of H_2_O or noni juice was placed.

*Single-fly two-choice assay*: 30-cell single-fly chambers were designed and manufactured by Formoplast S.A. following published blueprints (Gou et al., 2016), but using poly(methyl methacrylate) instead of acrylic, and adding a small handle to the top door. Flies were anesthetized in CO_2_ and placed in individual egg-laying chambers. Animals were allowed a 30 min period for recovery from anesthetization and acclimation to the chamber, during which time the oviposition substrate was prepared as described above. For single-odor assays, instant medium was dissolved in the desired odor solution diluted in water or juice. The concentration range of odors was defined from preliminary tests and previous studies (Amlou et al., 1998); specified concentrations represent those before adding the instant medium. For agarose substrates, 1 ml of agarose solution (containing the desired stimulus) was added to the bottom slot on one side of the oviposition chamber, which underlies 5 separate cells. For instant medium substrates, the paste was applied to the slot with a spatula. The fly loading slot of the single fly multi-chamber was placed on top of the bottom slot and flies accessed the instant medium. Eggs were scored on each substrate after 22 h. Preliminary tests of oviposition preferences on different substrates, which informed subsequent experimental design, are shown in Figure S2.

*Texture assays*: single-fly oviposition assays were performed as described above, but preparing substrates with different final concentrations of agarose.

For all assays, eggs were scored manually under a binocular microscope. The oviposition preference index was calculated as: (number of eggs in substrate X - number of eggs in substrate Y)/total number of eggs only when the number of eggs laid was ≥2. Indentations were scored as a small break/holes in the substrate surface; in some cases, the presence of multiple indentations in the same region of the substrate likely led to underestimation of the number of independent indentations.

### High-speed imaging of oviposition behavior

*D. sechellia* flies were cultured on standard cornmeal media. Mated *D. sechellia* females (aged 4-10 day-old in standard cornmeal food vials with or without supplementation with noni paste), were placed in empty vials with hydrated tissue for 14-20 h before the day of the experiment to force egg retention. A group of 2-3 individuals was introduced into a cubical oviposition filming chamber as described (Bracker et al., 2019). A trough on one side of the chamber was filled with a noni juice-agar substrate (1:3 or 1:6 v/v), with a wet pad at its base to limit sagging caused by desiccation. Flies were filmed with an external high-speed camera (JAI RMC-6740 GE, IMACO as detailed in (Bracker et al., 2019) using the custom FlyBehavior software (Bracker et al., 2019), focused on the surface of the agar column, which provided a 1 × 3 mm oviposition substrate. Recording of the group was performed for several separate 2 h-sessions over the course of one day (from late morning to mid-evening). The following specific behaviors were observed qualitatively by visual inspection of the resulting videos: *Touching* (Video S1) - the ovipositor simply touches the substrate; *Scratching* (Video S2) - the ovipositor brushes against the substrate, giving the impression of gentle scratching; *Digging* (Video S3) - the ovipositor burrows into the substrate surface; *Indentation formation* (Video S4) - the fly digs with its ovipositor at a particular site on the substrate leaving a minor depression (indentation) but no egg; occasionally, we observed that a fly returns to an indentation for egg-laying; *Egg laying* (Video S5) - the fly starts digging into the surface and lays an egg at this site. In many videos, we also observed flies exuding a liquid droplet, possibly from their anal plates (e.g., Video S1, example 4, left-hand animal); this action does not appear to be related to oviposition as it was observed also in virgin female and male flies (V.R. and N.G., unpublished). Video sequences were cropped and assembled in Fiji (Schindelin et al., 2012).

### Color preference assays

Color preference was assessed by adapting an olfactory trap assay (Prieto-Godino et al., 2017), in which the arena contained two traps filled with 300 μl of the same attractive odor – noni juice (*D. sechellia*) and balsamic vinegar (Antica Modena) (*D. melanogaster* and *D. simulans*) – masked with different visual cues. To simulate fruit at a ripe or unripe stage, a trap was covered with a green or white matte table-tennis ball (Lakikey; 40 mm diameter) with two opposing holes cut into it: one large hole on the bottom to insert the trap vial, and a smaller hole on top to allow the flies to enter the trap through a 200 μl pipette tip that was flush with the ball surface. Arenas were lined with either green or white paper (to provide different contrast for the green and white traps), and assays were performed in the light as well as under complete darkness in a behavior room (25°C and 60% relative humidity). Prior to the assay, flies were kept on standard media without noni supplement for 24 h. Twenty-five fed and mated 3-5 day-old females were introduced into each arena after brief ice anesthesia. The number of flies in each trap (as well as untrapped animals) was counted after 24 h; replicates where >25% of flies died within the experimental period were discarded. The preference index was calculated as: number of flies in white trap - number of flies in green trap/number living flies (trapped and untrapped).

### Locomotor activity monitoring

Activity was measured for 5-7 day old mated females at 25°C under a 12 h light: 12 h dark cycle, staged as for oviposition assays to ensure mating status, in the *Drosophila* activity monitor (DAM) system (Chiu et al., 2010) in incubators with continuous monitoring of light and temperature conditions (TriTech Research DT2- CIRC-TK). In brief, this system uses an infrared beam that bisects a 5 mm glass tube, in which the fly is housed, to record activity as the number of beam crosses per minute. Each tube is plugged with a 5% sucrose/2% agar (w/v) food source at one end and cotton wool at the other. Each DAM was used to record the activity of up to 32 flies simultaneously, and multiple monitors were contained in a single incubator. For all genotypes, we recorded flies over at least 2 technical replicates. Mean activity of an animal was calculated as the average number of beam crosses per minute over three complete days of recording.

### Ovary dissection and egg quantification

Females, prepared as for the oviposition assays, were anesthetized with CO_2_ and their ovaries dissected with forceps in phosphate buffered saline, using a surgical needle to separate the ovarioles. Mature eggs present in each ovary were counted under a binocular microscope.

### Statistical analyses

Oviposition preference indices were calculated compared to the null hypothesis (i.e., preference index = 0) for each strain using a Wilcoxon test with Bonferroni correction for multiple comparisons. Statistical differences across the number of eggs laid per fly per day for multiple comparisons were calculated applying Kruskal-Wallis rank sum test with Nemenyi post-hoc test. For two-sample comparisons, a two-sample t-test was used. The reference strain for multiple or two-sample comparisons is indicated in the figure legends. Error bars show SEM. All statistical values reported on the figures are as follows: NS (not significant) *P* > 0.05; * *P* < 0.05; ** *P* < 0.01; *** *P* < 0.001.

## Author contributions

R.A.-O. and R.B. conceived the project. R.A.-O performed all experiments except for those in Figures 3B, S3 and 6D, which were performed by M.P.S., and in Figure 2B and Videos S1-S5, which were performed by V.R., supervised by N.G. T.O.A. generated *D. sechellia* lines and contributed to the establishment of the assays in Figure 1B and 3E. All authors contributed to experimental design and interpretation of results. R.B. and R.A.-O. wrote the manuscript with contributions from all authors. All authors read and approved the final manuscript.

## Acknowledgements

We thank Chung-Hui Yang for valuable assistance in establishing the single-fly oviposition assay, Stefano Ceolin for advice on video recordings, René Gerber (University of Zurich Botanical Gardens) for a gift of *M. citrifolia*, Blaise Tissot-Dit-Sanfin for maintenance of *M. citrifolia*, Steeve Cruchet and Liliane Abuin for technical support, and members of the Benton laboratory for discussions and comments on the manuscript. V.R. was supported by the Graduate School of Systemic Neurosciences and by a Deutsche Forschungsgemeinschaft grant (GO 2495/9-1) to N.G. T.O.A. was supported by a Human Frontier Science Program Long-Term Fellowship (LT000461/2015-L) and a Swiss National Science Foundation Ambizione Grant (PZ00P3 185743). Research in R.B.’s laboratory is supported by the University of Lausanne, an ERC Advanced Grant (833548) and the Swiss National Science Foundation.

## Availability of data and materials

All materials and data supporting the findings of this study are available from the corresponding author on request.

---

**Figure S1.**
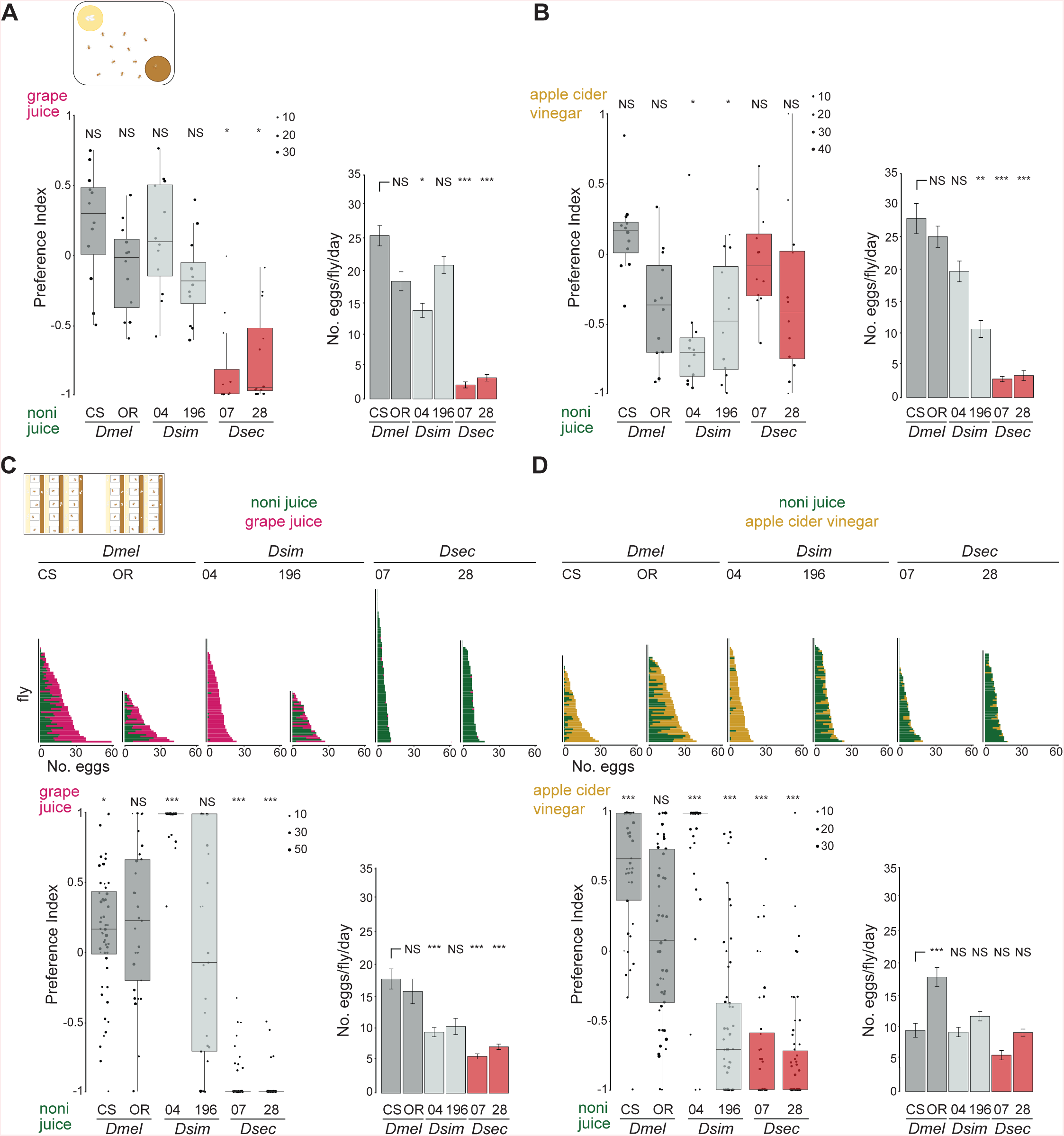
Oviposition preference assays using instant medium-based substrates. (A) Group oviposition preference assays for noni juice versus grape juice in instant medium 10g/100mL, using the same strains as in Figure 1C. Left: box plots of oviposition preference index. Statistical differences from 0 (no preference) are indicated (as in Figure 1C); n = 12 (representing 4 assays of 40-80 flies, each scored on 3 successive days with fresh oviposition plates each day). Right: bar plots of egg-laying rate. Statistically-significant differences from the *D. melanogaster* CS strain are indicated (as in Figure 1C). (B) Group oviposition preference assay, as in (A) for noni juice versus apple cider vinegar. n = 12 (representing 4 assays of 40-80 flies, each scored on 3 successive days with fresh oviposition plates each day). (C) Single-fly oviposition preference assay for noni juice versus grape juice in instant medium for the same strains as in (A). Top: total number of eggs laid in each substrate by each female. Bottom left: oviposition preference index. Bottom right: egg-laying rate. Mean values ± SEM are shown. (D) Single-fly oviposition preference assays, as in (C), for noni juice versus apple cider vinegar. n = 43-60 flies across 2 technical replicates.

**Figure S2.**
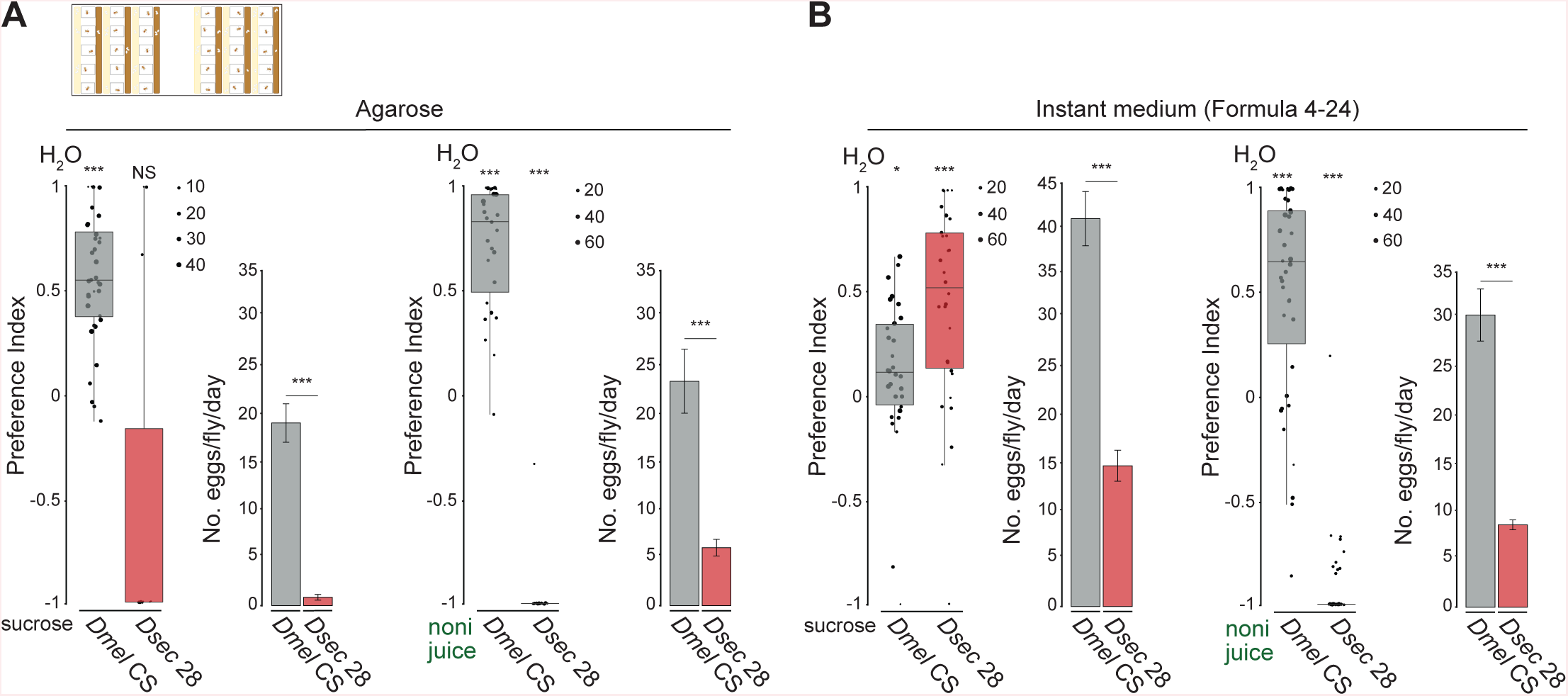
Establishment of single-fly oviposition assays. (A) Single-fly oviposition preference assays testing H_2_O versus either 150 mM sucrose or noni juice in agarose. Left: oviposition preference index. Statistical differences from 0 (no preference) are indicated: *** *P* < 0.001; * *P* < 0.05; NS *P* > 0.05 (Wilcoxon test with Bonferroni correction for multiple comparisons); n = 30-45 across 1-2 technical replicates. Preference of *D. melanogaster* for the plain the plain substrate resembles previous observations (Gou et al., 2016). To the right of each plot are bar plots of egg-laying rate. Mean values ± SEM are shown. *** *P* < 0.001 (two-sample t-test). (B) Single-fly oviposition preference assays, as in (A), in instant medium. n = 30- 45 across 1-2 technical replicates. The higher egg-laying rate of both species on instant medium compared to agarose substrates might reflect differences in texture and/or humidity; to avoid interfering with juices attraction, the vast majority of experiments were performed using agarose. Instant medium substrates were used in most single-odor experiments as egg-laying rate was very low on agarose substrates.

**Figure S3.**
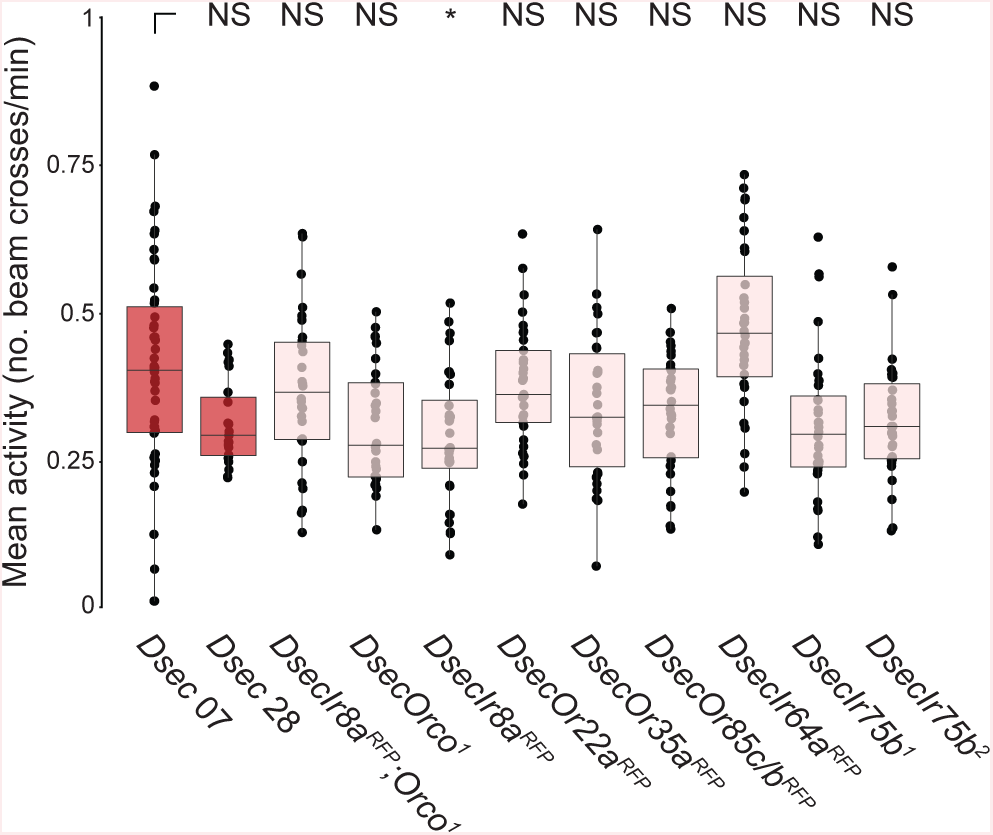
Locomotor activity levels of olfactory receptor mutants. Mean activity of *D. sechellia* mutant flies represented by the number of beam crosses per minute per fly recorded over three complete light-dark cycles. Statistical differences from *Dsec* 07 (the genetic background strain for all mutants) are indicated: * p < 0.05; NS *P* > 0.05 (Wilcoxon test with Bonferroni correction for multiple comparisons); n = 23-46 flies per genotype across at least 2 technical replicates.

**Figure S4.**
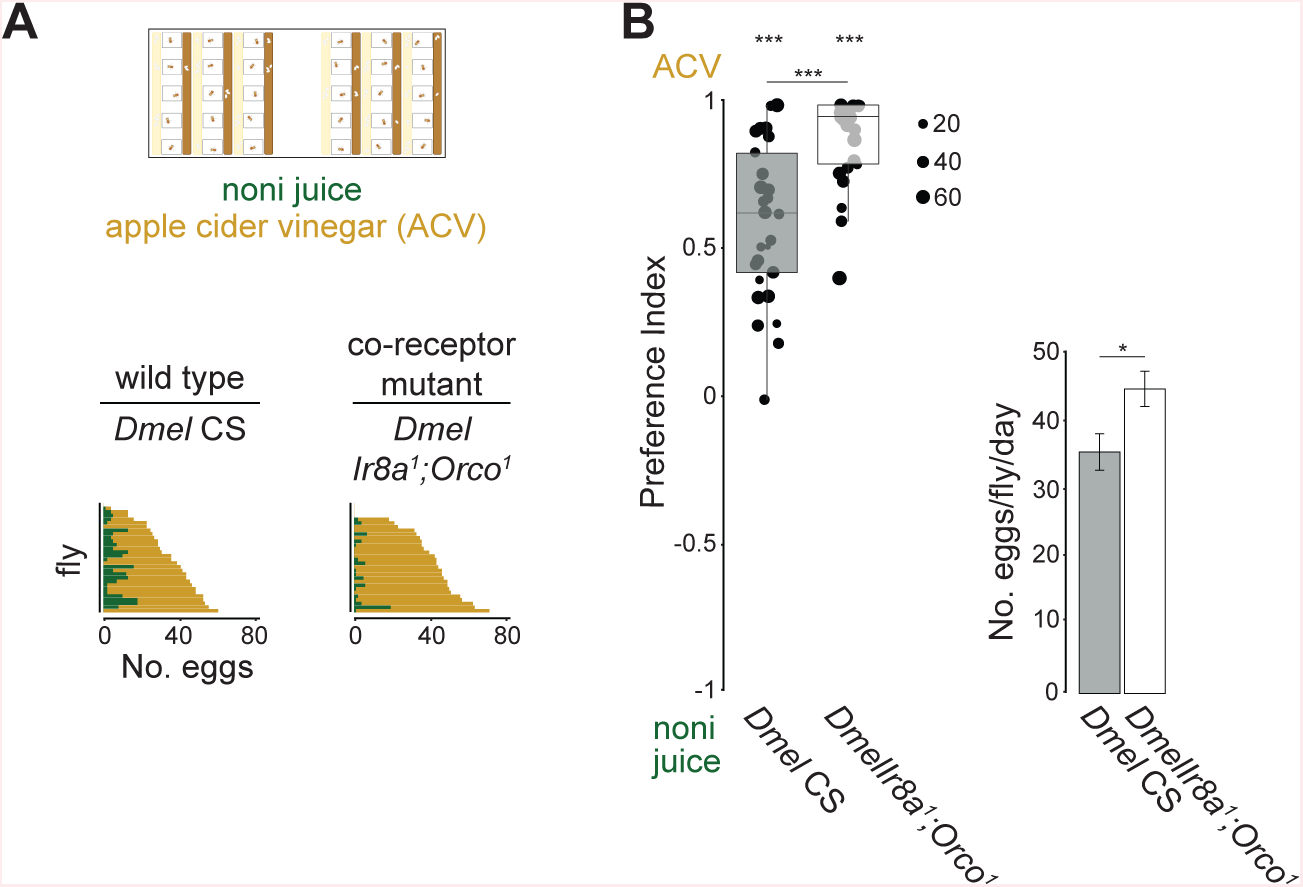
Near-anosmic *D. melanogaster* do not display defects in oviposition behavior. (A) Single-fly oviposition preference assays for apple cider vinegar versus noni juice in agarose for the indicated genotypes. The plots show the number of eggs laid per fly (n = 26-29 flies across 1 technical replicate). (B) Left: oviposition preference index for the assays shown in (A). Statistical differences from 0 (no preference) are indicated: *** *P* < 0.001 (Wilcoxon test with Bonferroni correction for multiple comparisons). *Dmel* CS and *DmelIr8a^1^/Orco^1^* show statistical difference (*P* = 6.334×10^-5^; Wilcoxon test with Bonferroni adjustment). Right: egg-laying rate. Mean values ± SEM are shown. Statistically-significant differences from the *D. melanogaster* CS strain are indicated: * *P <* 0.05 (Kruskal-Wallis rank sum test with Nemenyi post-hoc test). As the *Ir8a^1^;Orco^1^* mutant is not in the CS genetic background, it is unclear if the slight increase in preference for ACV and in egg-laying rate in the mutant strain reflects a role for these olfactory co-receptors or an effect of genetic background.

**Figure S5.**
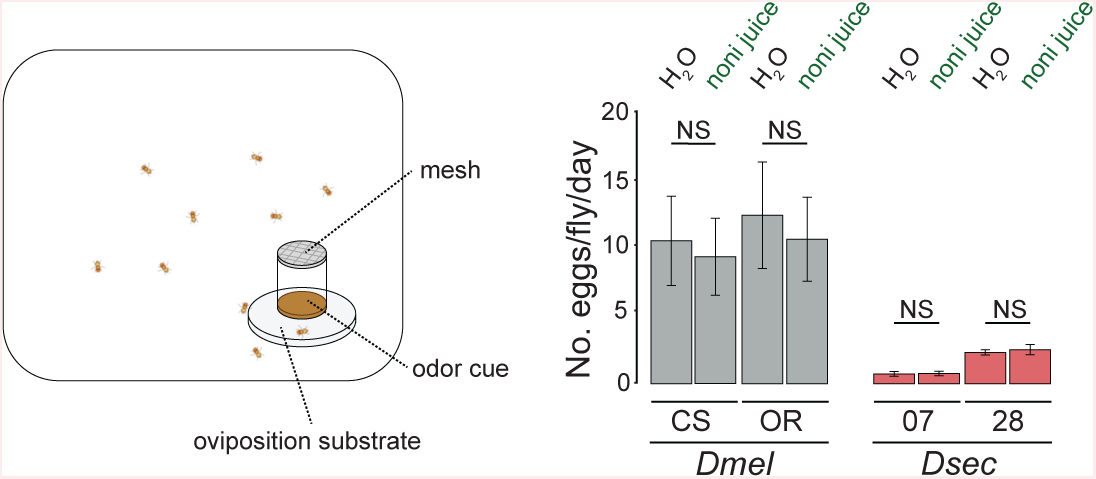
Olfactory cues are not sufficient for noni oviposition preference. Left: schematic of the one-choice group oviposition assay in the presence of a non-accessible noni juice source (or H_2_O control). Agarose plates supplemented with 150 mM sucrose were provided as oviposition substrates. Right: egg-laying rate. Mean values ± SEM are shown. NS *P* > 0.05 (Kruskal-Wallis rank sum test with Nemenyi post-hoc test); n = 12 (representing 4 assays, each scored on 3 successive days with fresh oviposition plates each day).

**Figure S6.**
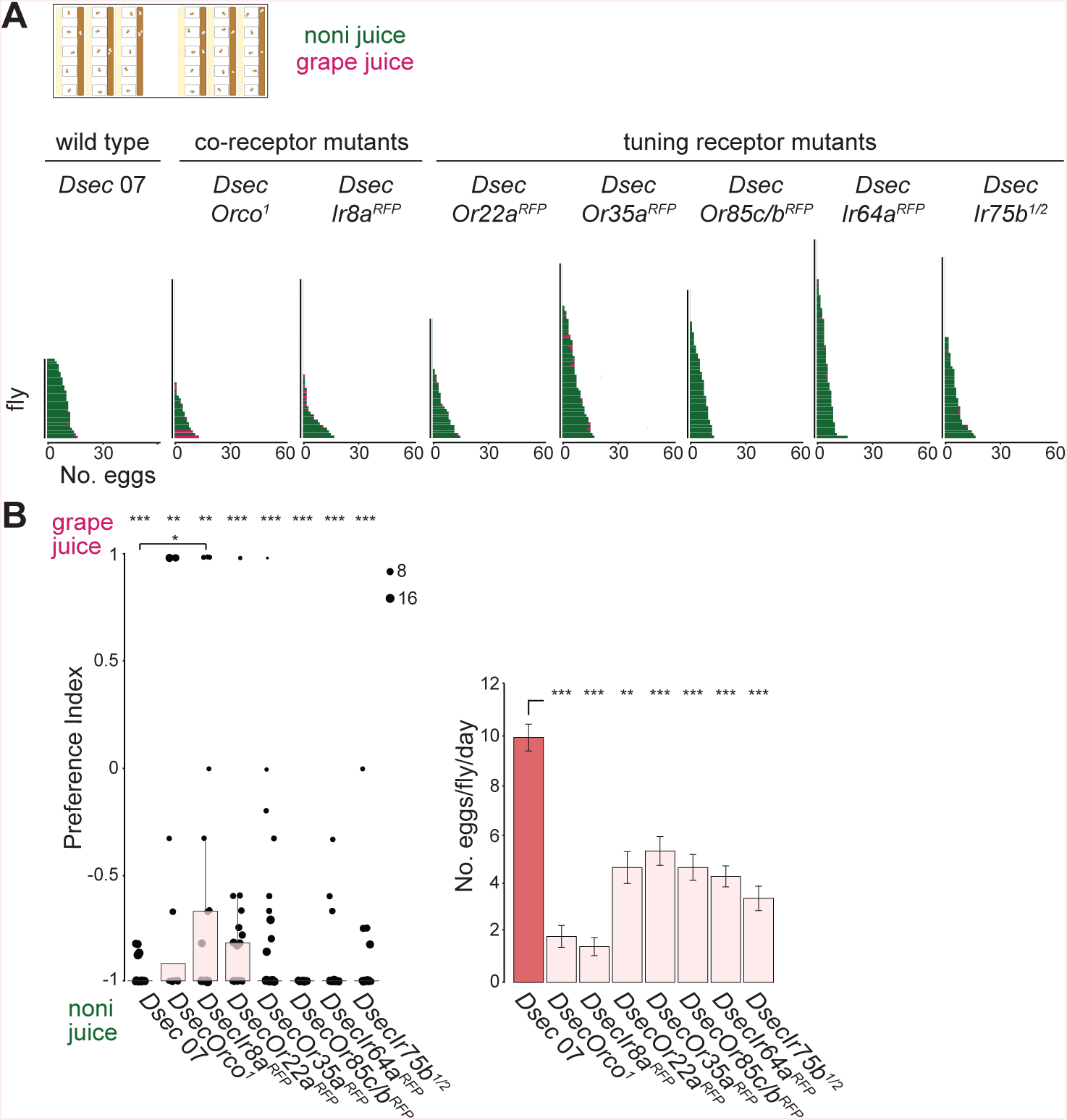
Olfactory pathways required for *D. sechellia* oviposition. (A) Single-fly oviposition preference assays for noni juice versus grape juice in agarose for the indicated genotypes. The plots show the number of eggs laid per fly (n = 30-60 flies across 1-2 technical replicates). (B) Single-fly oviposition assay for the same strains and conditions as in (A). Left: oviposition preference index. Statistical differences from 0 (no preference) are indicated: *** *P* < 0.001; ** *P* < 0.01; * *P* < 0.05 (Wilcoxon test with Bonferroni correction for multiple comparisons); n = 30-60 flies, 1-2 assays. *Dsec 07* and *DsecIr8a^RFP^* show statistical difference (*p*-value = 0.0223; Wilcoxon test with Bonferroni adjustment). Right: egg-laying rate in these assays. Mean values ± SEM are shown. *** *P* < 0.001; ** *P* < 0.01 (Kruskal-Wallis rank sum test with Nemenyi post-hoc test).

**Figure S7.**
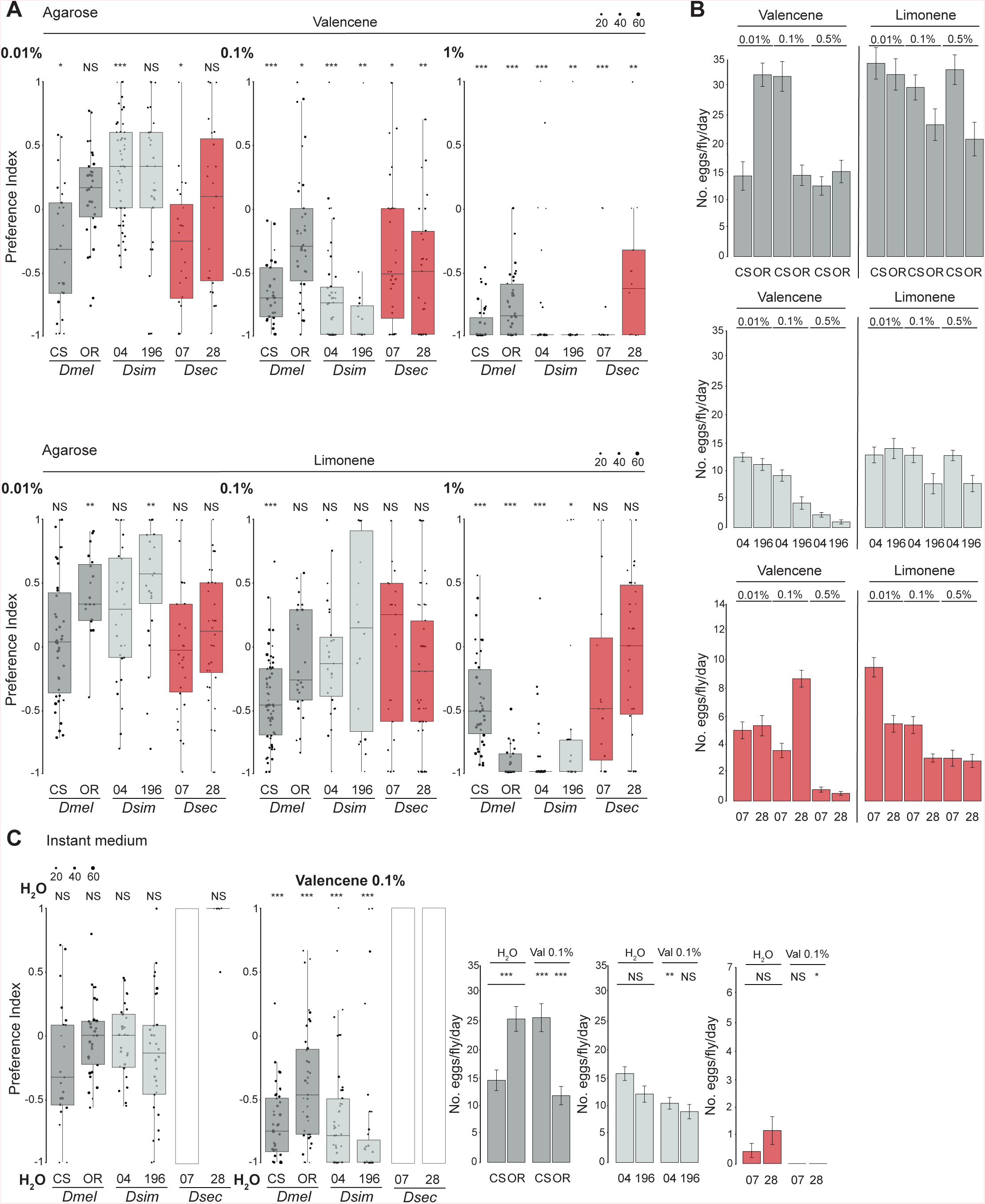
Influence of valencene and limonene on oviposition behavior. (A) Single-fly oviposition assays for valencene and limonene at the indicated concentrations. Odors were diluted in apple cider vinegar (for *D. melanogaster* and *D. simulans*) or noni juice (for *D. sechellia*) and agarose. For further information about fly strains and statistical meaning refer to the legend for Figure 1C. Statistical differences from 0 (no preference) are indicated: *** *P* < 0.001; ** *P* < 0.01; * *P* < 0.05; NS *P* > 0.05 (Wilcoxon test with Bonferroni correction for multiple comparisons); n = 30-90 flies, 1-3 assays. (B) Bar plots of egg laying rates for flies from (A). Mean values ± SEM are shown. (C) Single-fly oviposition assays of same strains as in (A), with an instant medium substrate. Left: oviposition preference index in control (H_2_O versus H_2_O) or experimental (0.1% valencene versus H_2_O) conditions. Statistical differences from 0 (no preference) are indicated: *** *P* < 0.001; NS *P* > 0.05 (Wilcoxon test with Bonferroni correction for multiple comparisons). n = 30-60 flies, 1-2 assays. Right: egg-laying rate in these assays. Mean values ± SEM are shown. Statistical comparisons of the effect of odors on egg-laying rate were performed across strains: *** *P* < 0.001; ** *P* < 0.01; * *P* < 0.05; NS *P* > 0.05 (two-sample t-test). For some strains (indicated with a white rectangle), the low number of flies laying eggs prevented calculation of a preference index.

**Figure S8.**
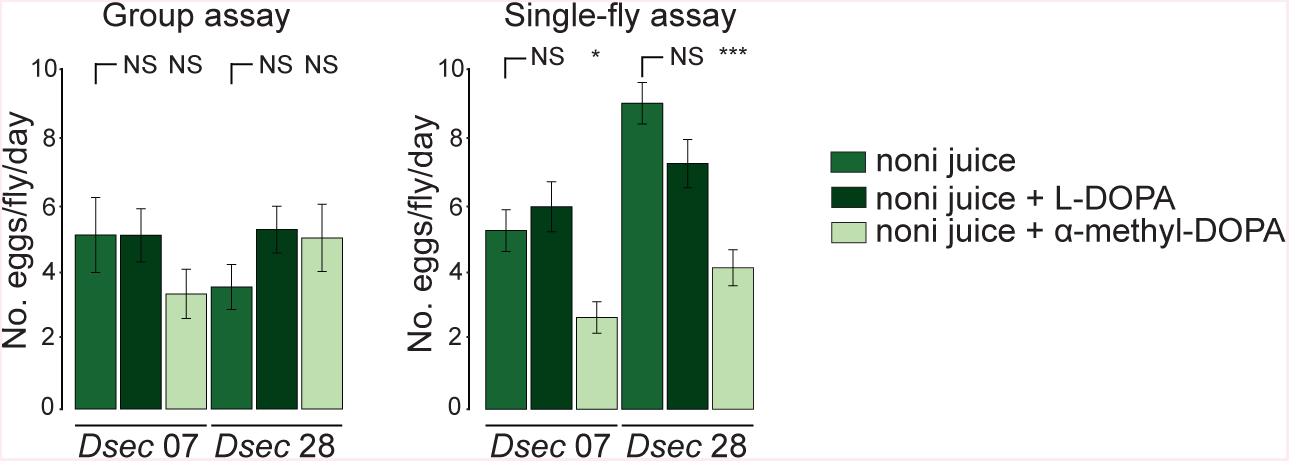
Impact of modulation of L-DOPA levels on egg-laying rate of *D. sechellia*. Egg-laying rate of *D. sechellia* cultivated on non-supplemented noni juice food media (green) or supplemented with L-DOPA or α-methyl-DOPA (see Methods). Left: group oviposition assays (representing 2 assays of either 10 (*D. melanogaster* and *D. simulans*) or 20 (*D. sechellia*) flies each, each scored on 3 successive days with fresh oviposition plates each day). Right: single-fly oviposition assays. Mean values ± SEM are shown (statistical differences from the noni juice substrates are shown: *** *P* < 0.001; * *P* < 0.05; NS *P* > 0.05 (Kruskal-Wallis rank sum test with Nemenyi post-hoc test); n = 60 flies across 2 technical replicates.

**Table S1.**
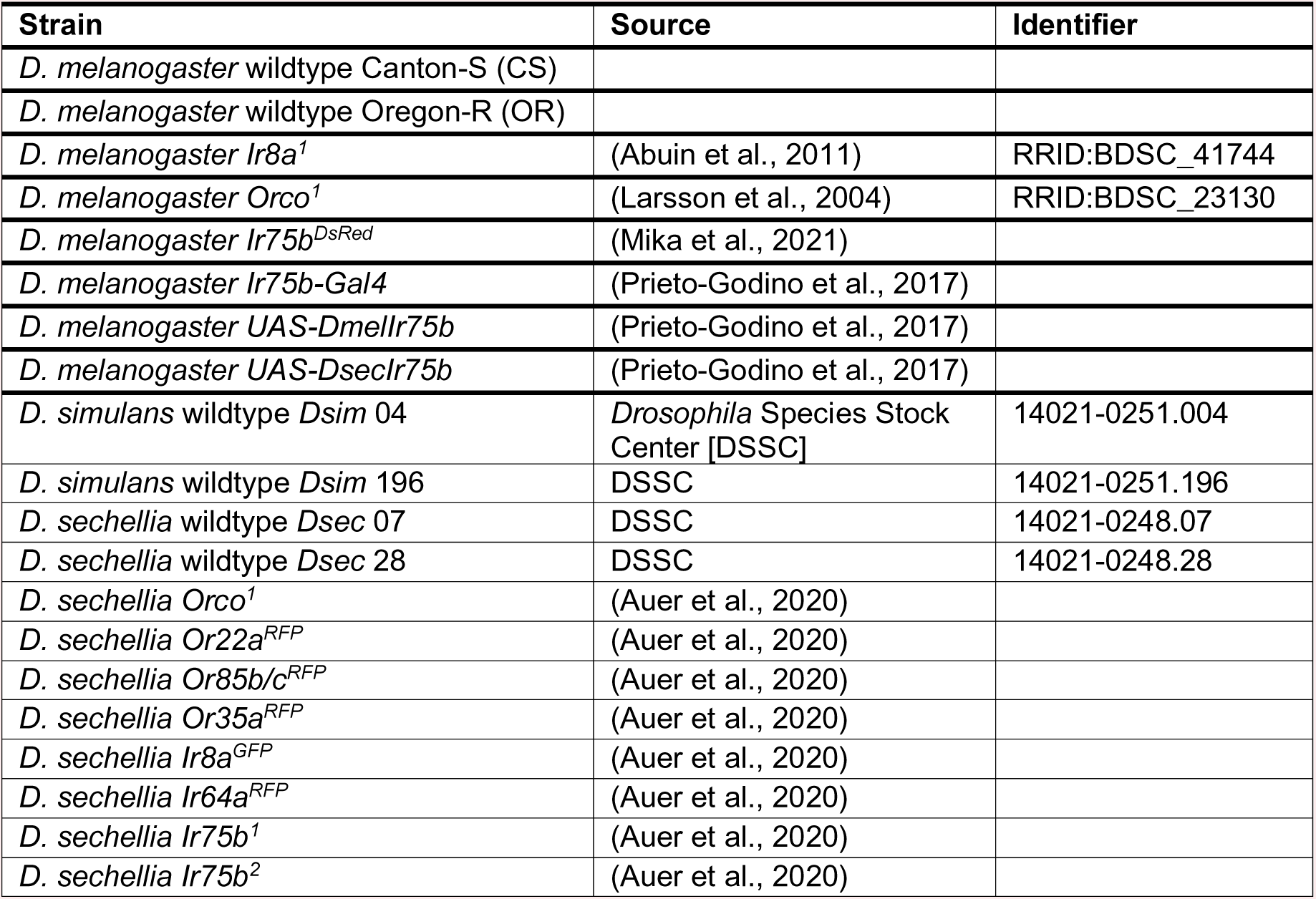
*Drosophila* strains.

**Table S2.** Chemicals.

| Odor | Supplier | CAS |
| --- | --- | --- |
| Agarose | Promega | - |
| Apple cider vinegar | Migros | - |
| BACTO Agar | BD | 214010 |
| Balsamic vinegar | Antica Modena | - |
| Butyric acid | Sigma-Aldrich | 107-92-6 |
| 3,4-dihydroxyphenylalanine (L-DOPA) | Sigma-Aldrich | 53587-29-4 |
| Formula 4-24® instant *Drosophila* medium ("instant medium") | Carolina Biological Supply | - |
| Grape juice | Beutelsbacher Bio | - |
| 2-heptanone | Sigma-Aldrich | 110-43-0 |
| Hexanoic acid | Sigma-Aldrich | 142-62-1 |
| Limonene | Sigma-Aldrich | 5989-27-5 |
| α-methyl-DOPA | Sigma-Aldrich | 41372-08-1 |
| Methyl hexanoate | Sigma-Aldrich | 106-70-7 |
| Noni juice | Raab Vitalfood Bio | - |
| Octanoic acid | Sigma-Aldrich | 124-07-2 |
| Sucrose | Sigma | 57-50-1 |
| Valencene | Sigma-Aldrich | 4630-07-3 |

**Video S1. Compilation of video sequences illustrating substrate touching by the ovipositor in *D. sechellia*.**

**Video S2. Compilation of video sequences illustrating substrate scratching by the ovipositor in *D. sechellia*.**

**Video S3. Compilation of video sequences illustrating substrate digging by the ovipositor in *D. sechellia*.**

**Video S4. Compilation of video sequences illustrating indentation formation by the ovipositor in *D. sechellia*.**

**Video S5. Compilation of video sequences illustrating egg-laying by *D. sechellia*.**

